# Assessing coverage of the Monitoring Framework of the Kunming-Montreal Global Biodiversity Framework and opportunities to fill gaps

**DOI:** 10.1101/2024.09.25.614896

**Authors:** F. Affinito, S. H. M. Butchart, E. Nicholson, T. Hirsch, J. M. Williams, J. Campbell, M. F. Ferrari, M. Gabay, L. Gorini, B. Kalamujic Stroil, R. Kohsaka, B. Painter, J. C. Pinto, A. H. Scholz, T. R. A. Straza, N. Tshidada, S. Vallecillo, S. Widdicombe, A. Gonzalez

## Abstract

The Kunming-Montreal Global Biodiversity Framework (GBF) is the most ambitious agreement on biodiversity conservation and sustainable use to date. It calls for a whole-of government and whole-of-society approach to halt and reverse biodiversity loss worldwide. The Monitoring Framework of the GBF lays out how Parties to the Convention on Biological Diversity (CBD) are expected to report their progress. A CBD expert group provided guidance on its implementation, including a gap analysis to identify the strengths and limitations of the indicators in the Monitoring Framework. We present the results of the gap analysis, highlight where more work is needed and provide recommendations on implementing and improving monitoring to allow effective and comprehensive tracking of the GBF’s ambition. We find that with the headline and binary indicators, which Parties are required to use, the Monitoring Framework fully covers 19% of the GBF’s ambition and partially covers an additional 40%. Including disaggregations of the headline indicators improves coverage to 22% fully and an additional 41% partially. Adding optional (component and complementary) indicators brings full coverage to 29% with an additional 47% partial coverage. No indicators are available for 12% of the GBF. In practice, the coverage of the Monitoring Framework will depend on which indicators (headline and binary as well as component and complementary) and disaggregations are used by countries. Disaggregations are particularly relevant to monitor the cross-cutting considerations defined under section C. Substantial investment is required to collect the necessary data to compute indicators, infer change, and effectively monitor progress. We highlight important next steps to progressively improve the efficacy of the Monitoring Framework.

## 1. Introduction

Since the Convention on Biological Diversity (CBD) was opened for signature at the United Nations Conference on Environment and Development (also known as the ‘Earth Summit’) in 1992^1^, three sequential decadal plans have been adopted to address the alarming rate of biodiversity loss worldwide^2^. The targets for 2010 and 2020 were largely unmet^3–5^. The latest plan, the Kunming-Montreal Global Biodiversity Framework (GBF), adopted in 2022, is the most ambitious to date^6^. The GBF hinges upon a 2050 vision of living in harmony with nature, to be realized through commitments towards four Goals by 2050 and 23 action-oriented Targets by 2030^6^. The GBF is built around a theory of change (Figure 1) that aims to tackle the drivers of biodiversity loss, recognising that urgent, transformative, and widespread action is required across all sectors of society to halt and reverse biodiversity loss.

**Figure 1.**
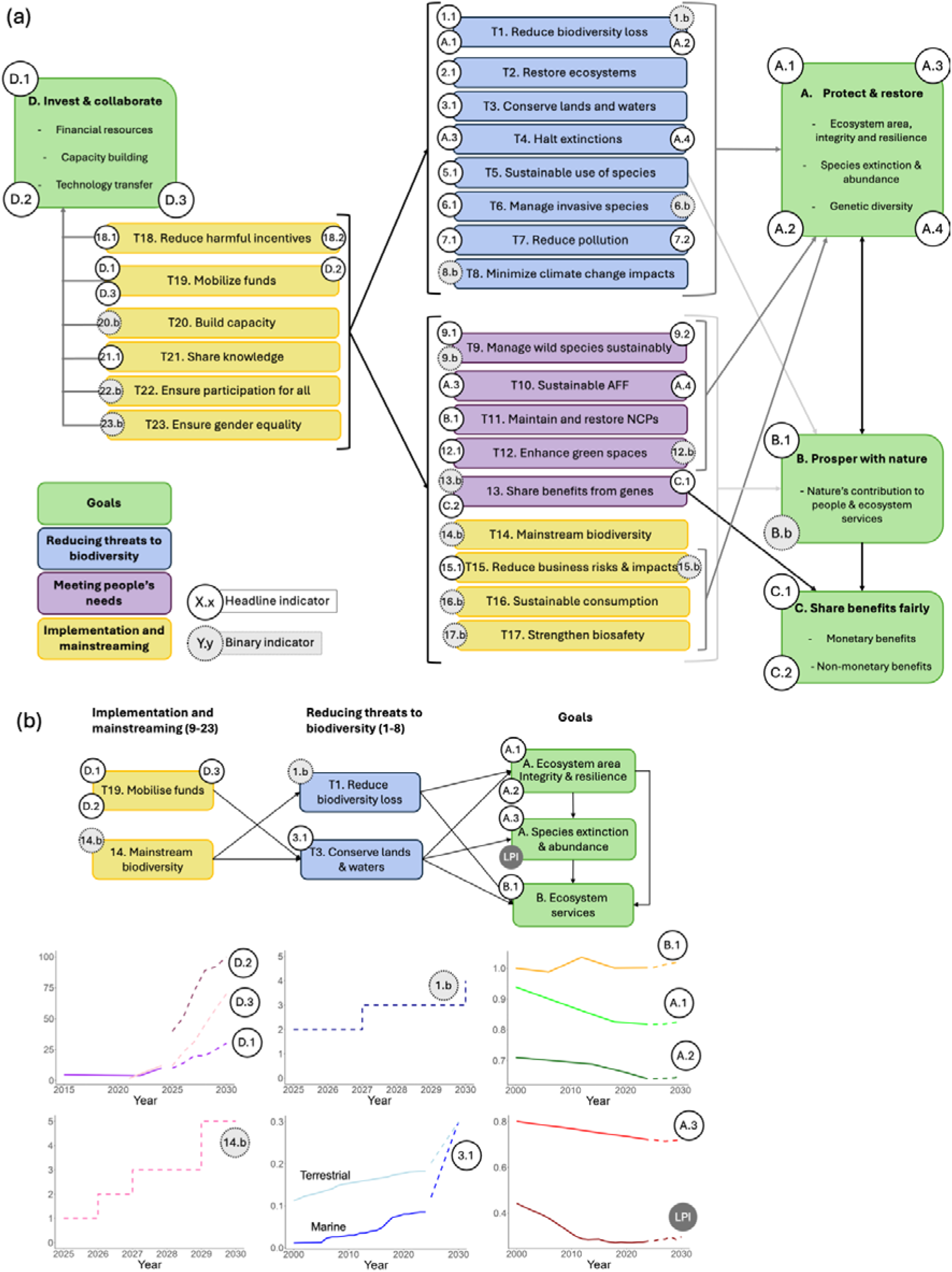
The GBF theory of change organised by Goals and Targets with their required indicators. (a) The GBF divides its Targets into three categories: reducing threats to biodiversity (Targets 1 to 8), meeting people’s needs (Targets 9 to 13), and implementation and mainstreaming (Targets 14 to 23), that are needed to deliver on its ambition and achieve the four Goals. All of them have at least one required indicator (headline, binary or both, in circles) to track progress and some of them are related. Implementation of the Targets by 2030 contributes to achieving the Goals and other linked Targets in the GBF. The indicators in the Monitoring Framework are designed to help track progress along all aspects of the GBF. (b) A partial theory of change for some Targets (1, 3, 14 and 19) and Goals (A and B) of the GBF, with associated indicators, showing how mainstreaming and investment can increase protection and conservation, leading to the Goals for species, ecosystems and ecosystem services. Reporting systematically on the required indicators will allow progress to be tracked nationally and globally on each Goal and Target, enabling action to be taken as needed to deliver on the overall theory of change. Parties may also choose to report on optional indicators (e.g. the LPI – Living Planet Index). Quantitative headline and component indicators can track yearly changes whilst binary indicators are reported with every national report. Where past data is available, graphs show solid lines. Dashed lines represent optimistic predictions of a world where Goals and Targets are met. Indicators that are new are shown only with dashed lines as no past data is available to estimate them. AFF stands for agriculture, forestry and fisheries; NCP stands for nature’s contributions to people.

The GBF includes a Monitoring Framework (Box 1), an important innovation guiding countries on how to report their progress towards the Goals and Targets (Figure 1). The Monitoring Framework sets out how Parties are expected to record their efforts and progress using a set of consistent indicators compiled at the national level^7,8^, while allowing some flexibility due to differing capacity and data availability across Parties^9^. The Monitoring Framework was established to improve transparency and accountability among Parties^10^, recognising that failure to reach targets has been linked to low levels of implementation^11^ and difficulties in tracking progress^12^. Parties are requested to use the Monitoring Framework in future national reports to the CBD, including the next (seventh) national report due in February 2026, ahead of the 17^th^ meeting of the Conference of the Parties (COP17). There is now a need for the scientific community and relevant actors to support countries with the design and implementation of their domestic monitoring frameworks, including how monitoring programs can gather the data needed for indicator updates^13^.

### Box 1. The structure of the Monitoring Framework

#### Indicators

The Monitoring Framework relies on five types of indicators designed to track progress towards the Goals and Targets of the GBF^7^. In the context of the CBD, indicators can be defined as “a measure based on verifiable data that conveys information about more than itself”^133^. These indicators include headline and binary indicators (which Parties are required, and expected, to report on), component and complementary indicators (which are optional), and additional nationally relevant indicators that Parties may choose. Through the structure provided by its indicators, the Monitoring Framework enables much greater standardisation of the outputs expected from Parties in their national reports, compared with previous reporting rounds, marking a paradigm shift.

Headline indicators are a minimum set of high-level indicators intended to capture the overall scope of a Goal or Target. There are a total of 26 headline indicators in the framework spread across the four Goals and 15 of the 23 Targets. Headline indicators are quantitative measures of process and/or outcomes relevant to each Goal or Target. These indicators are calculated from data mostly collated at the national level but can be aggregated or disaggregated to provide more in-depth information - for example by taxonomic group for species-related indicators or by sectors of society (e.g. gender and youth). In some cases, these data are obtained from third-party multilateral data aggregation tools, such as the World Database on Protected Areas. Headline indicators each have between 2 and 21 disaggregations recommended by the AHTEG and described in the metadata for each indicator. The use of recommended disaggregations is optional.

Binary indicators are qualitative measures of the efforts made by Parties to deliver on the GBF. These indicators are compiled from a set of questions to be answered by Parties and aimed at understanding the progress made towards implementing measures, processes, and legislation to deliver on the ambition of the GBF^116^. The answers to the set of questions can be used to assign an overall score (from 0 to 5, showing an increasing level of implementation) for each binary indicator, providing valuable information about the progress made by Parties in facilitating and promoting the outcomes of the Goals and Targets. These indicators are simpler in design than headline indicators and are more readily compiled, though detailed consideration by Parties on which answer to choose will be needed.

Component and complementary indicators help monitor progress made towards the Goals and Targets beyond the information provided by the headline and binary indicators. Currently, 55 component and 124 complementary indicators are spread across all of the Goals and Targets^130^. Component indicators are intended to cover elements of the Goals and Targets that are not covered by headline or binary indicators. Complementary indicators support a more in-depth thematic analysis of the Goals and Targets.

Finally, national indicators include any additional indicators that Parties have established or developed, which they may choose to compile and report on, supplementing the other indicators. These national indicators may be relevant to the specific context of a Party and provide additional information on progress.

##### National reports

Regular national reports presented by Parties provide information on measures taken in support of the Convention’s objectives and their outcomes. Six rounds of reporting were completed between 1994 and 2020; the seventh and eighth national reports are due in 2026 and 2029 respectively (Article 26^134^). In the upcoming 7^th^ and 8^th^ national reports, Parties will be required to report on the headline indicators using data assembled through biodiversity and ecosystem monitoring networks and relevant agencies and to provide responses for all the binary indicators. Where indicators are generated by international bodies using local, national, regional, and/or global data, the values for each country will be pre-populated in national report templates. Parties can choose to use these values or replace them with their own where appropriate. Parties may further choose to include any additional optional indicators. Using the information submitted in national reports through the online reporting tool^135^, the CBD Secretariat will be able to compile and track progress towards the GBF at the global level, for example in the global report^136^ on collective progress in the implementation of the GBF, due in 2026.

The effectiveness of the GBF’s Monitoring Framework depends on three key aspects: 1) how well its indicators cover the scope of the GBF and its Goals and Targets; 2) national uptake of the Monitoring Framework as a driver for improving national monitoring systems, including the standardized methodologies; and 3) the dissemination and sharing of data and metadata on the indicators of the Monitoring Framework, including in the national reports to the Convention. Despite extensive research and use of scientific data and evidence in monitoring biodiversity and ecosystem processes^14–17^, significant gaps remain, including in geographic and taxonomic data coverage for some indicators^18–23^, and a lack of relevant and available indicators for areas such as biodiversity-relevant measures of human wellbeing^24^. To address these challenges, the Monitoring Framework’s success will depend upon mobilizing national, regional and international actors to support data collection and indicator compilation.

The CBD’s Subsidiary Body on Scientific Technical and Technological Advice (SBSTTA) requested a monitoring framework be developed in 2019^25^. The early processes of designing the Monitoring Framework occurred alongside the negotiation of the GBF text and focused on selecting indicators from available lists, such as the Sustainable Development Goals indicators^26^ and those generated by members of the Biodiversity Indicators Partnership. This resulted in a list of indicators in various stages of development that was then categorized and matched with relevant Goals and Targets. The selection process called for at least one headline indicator (Box 1) per Goal and Target. Where possible, headline indicators were assigned to quantitatively assess progress towards the main intent of the Goal or Target. Headline indicators were intended to comprise a small list of indicators to capture the overall scope of a Goal or Target. This *post hoc* process resulted in some Targets and Goals lacking headline indicators. Binary indicators (Box 1) were developed to qualitatively assess progress towards a Goal or Target (for example, does a legislative framework exist to address a Target’s aims). Where existing indicators were deemed useful for covering a single aspect of a Goal or a Target, but less relevant for covering its full scope, these were assigned to the optional component or complementary indicator list (Box 1) to allow Parties wishing to report on progress in more detail to do so. Additionally, indicators which had limited geographic coverage or which may not be useful or possible to implement for most Parties were assigned to the optional component or complementary indicator list (Box 1). The list of indicators in the Monitoring Framework represents a political agreement from Parties on the aspects of GBF which were determined to be the basis for monitoring its implementation.

The scientific guidance for operationalising the Monitoring Framework followed the political process of selecting indicators^8^. Parties established an Ad Hoc Technical Expert Group (AHTEG) on Indicators in April 2023 to provide guidance for operationalising the Monitoring Framework ahead of the COP16 meeting in October 2024. This expert group comprised 45 experts nominated by Parties and observer organisations^27^ and was tasked with reviewing all the indicators, producing methodological guidance for implementing the Monitoring Framework, and assessing how well it covers the ambition of the GBF^28^. The AHTEG included individuals proposed by Parties and observer groups, selected for their expertise on the different aspects of the GBF and representing a diversity of scientific, technical, and policy fields as well as Indigenous Peoples and Local Communities (IPLCs) and Youth.

The AHTEG worked with headline indicator developers and agencies supporting methodological research and capacity building to ensure that each of these indicators had a robust methodology while considering the real capacity constraints, limited data availability, and other challenges faced by many Parties (Figure 2). The AHTEG also reviewed all component and complementary indicators to determine the availability of their methodology. The AHTEG conducted a gap analysis, through a multi-step expert elicitation process^29–32^ (Methods), to assess the degree to which the current indicators (headline indicators, their recommended disaggregations, and binary, component, and complementary indicators) cover all substantive elements of the Goals and Targets. This gap analysis aimed to highlight limitations in the Monitoring Framework and support Parties in identifying opportunities to improve its effectiveness and coverage, given that Parties had agreed to keep the Monitoring Framework under review^7^.

**Figure 2.**
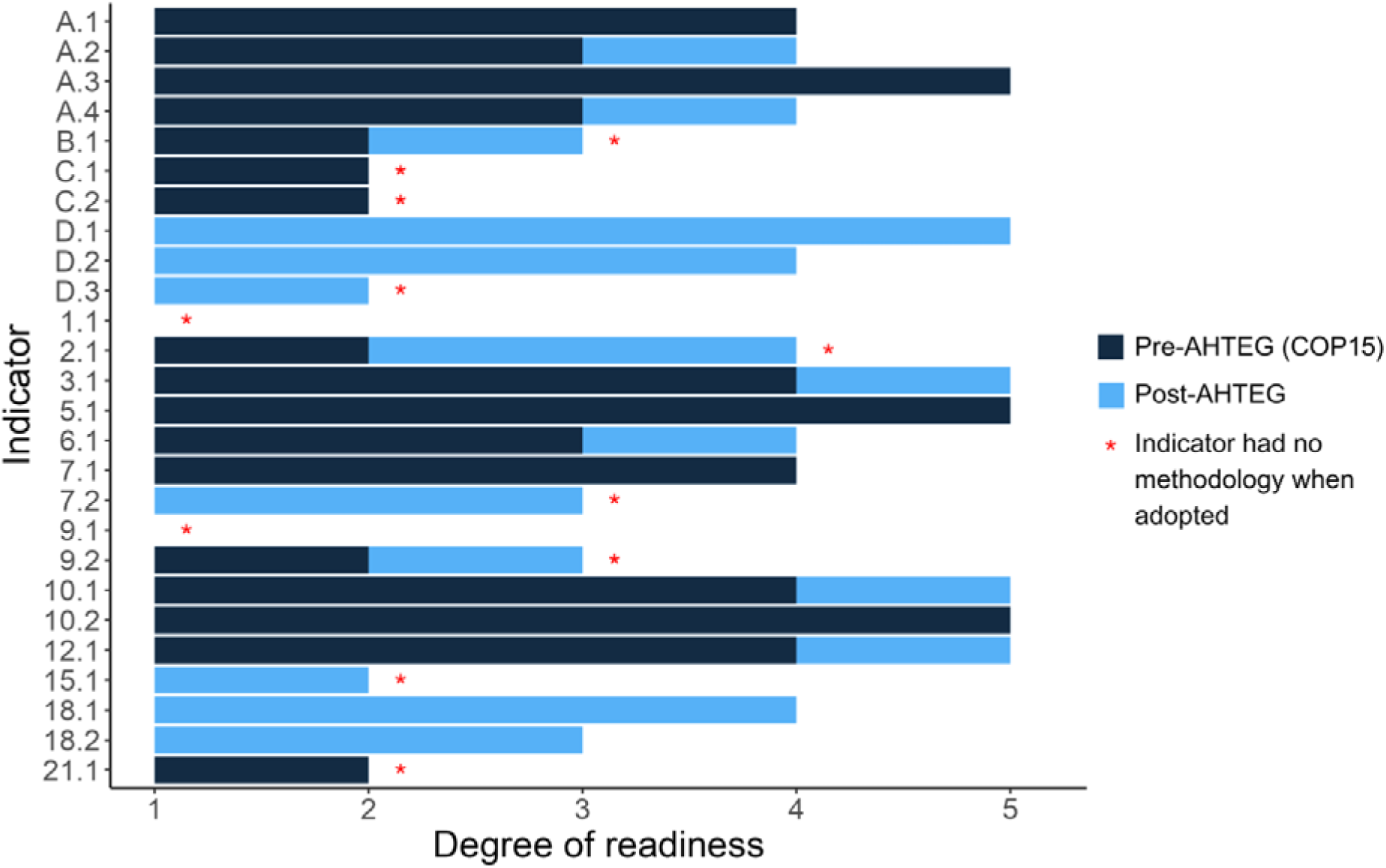
Degree of readiness of the headline indicators for the Global Biodiversity Framework. Dark blue bars show the readiness level at the time of COP15 (December 2022), and light blue bars show the level in June 2024 (i.e. reflecting progress since COP15). Asterisks identify indicators that lacked an agreed methodology when they were adopted at COP15^7^. Readiness was scored as: 1. Methods not yet developed, and a process needs to be established to develop them. 2. Methods not yet developed but a process is underway, led by one or more organisations, to develop them. 3. Methods developed (or partially developed) and tested/piloted, but data not yet widely available (and/or collection not yet underway). 4. Methods established, data being compiled and indicator operational in at least some countries, but further investment in methods ongoing and/or further data collection required. 5. Methods established, data being compiled and accessible, and indicator operational for most/all countries.

To assess the Monitoring Framework’s coverage, the text of each Goal and Target was split into elements (190 total) that reflect distinct, independently measurable objectives that each need to be achieved for full implementation of the GBF (Methods). For each element, indicators were evaluated and their coverage was assessed (see Box 2 for an example, Supplementary Table 2 for the complete list of elements). The AHTEG identified gaps and made general recommendations to Parties on improving the Monitoring Framework^33^. In this article, we – members of the AHTEG and collaborators in the process – (i) present the results of the gap analysis and review of the cross-cutting issues of the Monitoring Framework, (ii) share recommendations made to Parties and (iii) suggest next steps for effectively monitoring the implementation of the GBF.

### Box 2. Gaps in indicators for Target 12

Here, an example for Target 12 shows the separation of the target into its individual elements and coverage by indicators. The specific emphasis of each element to be monitored in the Monitoring Framework is highlighted in **bold**. This target contains eight (a–i) distinct elements that reflect its ambition to transform the relationship between cities and biodiversity. All other targets were similarly assessed in the gap analysis. Target 12 has a headline (12.1), binary (12.b), and component (City Biodiversity Index – CBI^3^) indicator. Headline 12.1 was adopted by Parties in part because it is an SDG indicator with a development team and host organisation based at UN-Habitat. This headline indicator however falls short of measuring change in most of the elements included in Target 12 because it only quantifies access to green spaces, which partially informs on element 12e (access to green and blue spaces) and could provide some indirect information on 12a (area of green spaces). Indicator 12.b further tracks the existence of measures in place for biodiversity-inclusive urban planning, enabling monitoring of 12g. The other elements of Target 12 are not covered by its headline or binary but some (12a, b, f, and h) could be monitored for those Parties choosing to report on the component indicator. Overall, monitoring progress on Target 12 under the Monitoring Framework is therefore only currently guaranteed for one element, partially feasible for one element, potentially feasible and dependent on Parties’ good will for four elements, and not possible for two elements. In this case, the component indicator for Target 12 appears theoretically superior to its headline indicator but is not supported by any organisation and requires a large amount of data which is unlikely to be available in most cities. It is worth noting that the City Biodiversity Index was not a component indicator in the originally negotiated Monitoring Framework but was recommended as such by the AHTEG.

| **Target 12:** Significantly increase the area and quality and connectivity of, access to, and benefits from green and blue spaces in urban and densely populated areas sustainably, by mainstreaming the conservation and sustainable use of biodiversity, and ensure biodiversity-inclusive urban planning, enhancing native biodiversity, ecological connectivity and integrity, and improving human health and well-being and connection to nature and contributing to inclusive and sustainable urbanisation and the provision of ecosystem functions and services. |  |  |  |
| --- | --- | --- | --- |
| Target 12 elements | Mandatory indicator(s) | Coverage | Optional indicator(s) |
| 12a. Significantly increase the **area of green spaces in urban and densely populated areas** | 12.1<sup>1</sup> | Potential | CBI<sup>3</sup> |
| 12b. Significantly increase the **area of blue spaces in urban and densely populated areas** |  |  | CBI<sup>3</sup> |
| 12c. Significantly increase the **quality of green and blue spaces in urban and densely populated areas** |  |  |  |
| 12d. Significantly increase the **connectivity of green and blue spaces in urban and densely populated areas** |  |  |  |
| 12e. Significantly increase the **access to and benefits from green and blue spaces in urban and densely populated areas** | 12.1<sup>1</sup> | Partial |  |
| 12f. **Mainstream the conservation and sustainable use of biodiversity in urban and densely populated areas** |  |  | CBI<sup>3</sup> |
| 12g. Ensure **biodiversity-inclusive urban planning** contributing to inclusive and sustainable urbanization and to the provision of ecosystem functions and services. | 12.b<sup>2</sup> | Complete |  |
| 12h. **Enhance native biodiversity in urban and densely populated areas** |  |  | CBI<sup>3</sup> |
| 12i. **Improve human health and well-being and connectedness to nature,** through biodiversity-inclusive urban planning |  |  |  |

<sup>1</sup> Headline indicator 12.1 is based on SDG indicator 11.7.1 which measures the share of land allocated to public spaces and the total population with access to these spaces by age, gender, and disability.
<sup>2</sup> Binary indicator 12.b tracks the number of countries with biodiversity-inclusive urban planning referring to green or blue urban spaces.
<sup>3</sup> The City Biodiversity Index is built on a score of 28 indicators measuring different aspects of a city's biodiversity and relationship to biodiversity <sup>45</sup>.

## 2. Gap analysis

### 2.1 Overall coverage

The indicators of the Monitoring Framework that are required in the national reports (headline without disaggregations and binary) fully cover 19% (36/190) of the elements of the Goals and Targets and partially cover an additional 40% (76/190). Applying the recommended disaggregations of headline indicators increases this coverage to 22% (42/190) fully and an additional 41% (78/190) partially. The use of the indicators that are optional in the national reports (component and complementary) further broadens the coverage of the Monitoring Framework to 29% (55/190) fully and an additional 47% (90/190) partially (Figure 3). Additionally, the use of recommended disaggregations of headline indicators, combined with the optional component and complementary indicators, reduces the number of elements excluded from monitoring of the Goals and Targets from 29% (56/190) to 12% (23/190), which implies a reduction of 59% in the number of gaps in the Monitoring Framework.

**Figure 3.**
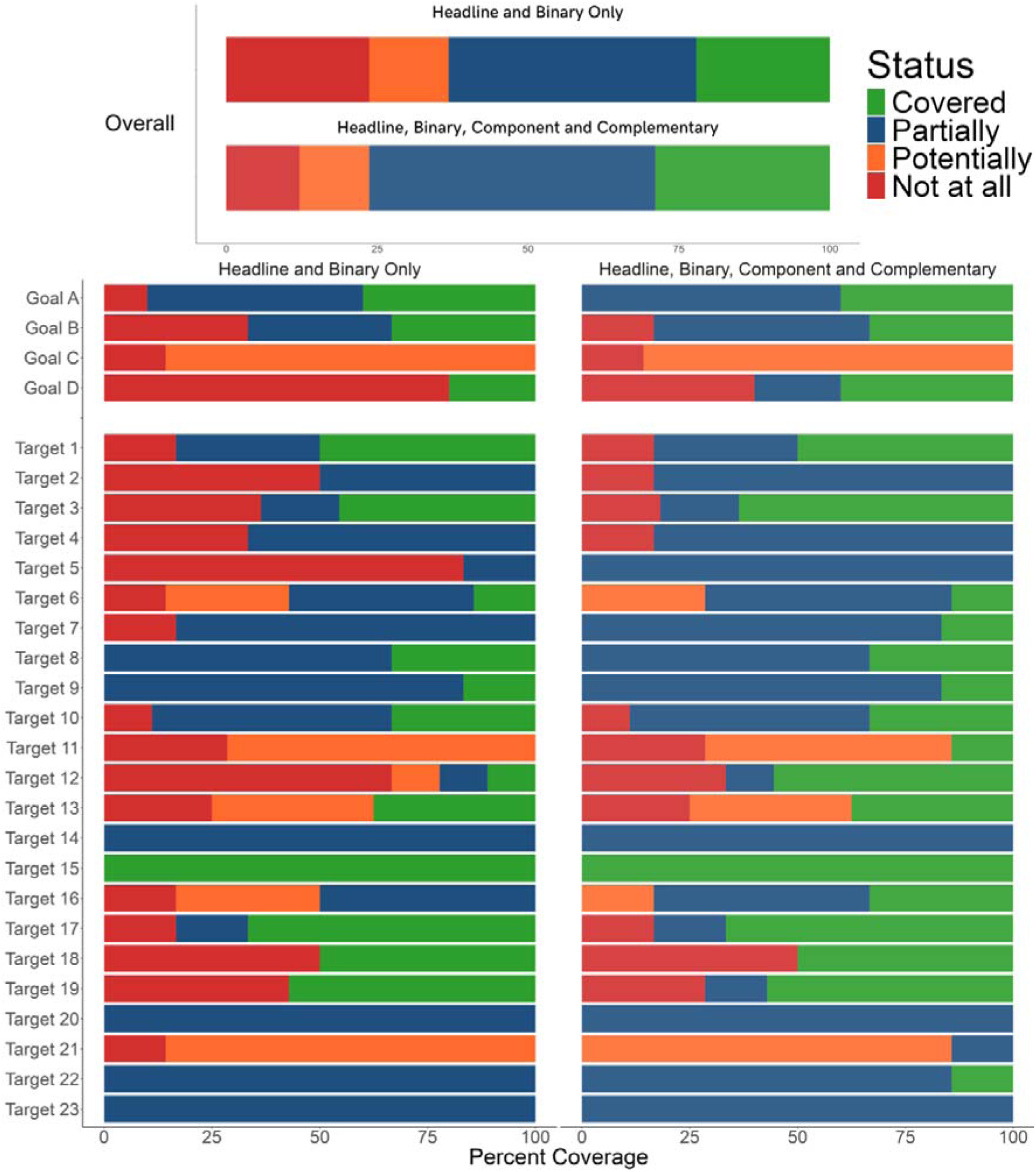
Coverage of elements of the Goals and Targets of the GBF by the indicators in the Monitoring Framework. On top, overall coverage of the indicators for all Goals and Targets in the GBF. On the left, coverage by the required headline and binary indicators for each Goal and Target (including the recommended disaggregations of the former). On the right, coverage by these indicators as well as the optional component and complementary indicators for each Goal and Target. ‘Partially covered’ applies to elements for which the indicator(s) track progress towards some aspects of the element, but not all. ‘Potentially covered’ applies to elements that could be covered by indicators that are still in development, so there is uncertainty as to whether the final metric(s) produced will adequately cover the element. ‘Not at all’ means that no indicators (headline, binary, component, nor complementary) are available to monitor the element.

Importantly, many elements are not currently covered (*i.e.* classed as gaps or potentially covered) by the indicators in the Monitoring Framework: 36-50% for Goals (10-14/28, depending on whether recommended disaggregations, and optional component and complementary indicators are included) and 25-40% for Targets (35-64/162). Ranges show values with and without inclusion of all disaggregations, component and complementary indicators. If all Parties report only on the required indicators coverage will be limited to the lowest range value; coverage will be even lower if they are unable to report on some required indicators (e.g. if data are not available). The highest range value will be reached if all Parties additionally report on all component and complementary indicators. Realistically, we can expect some coverage in between to be achieved as some Parties choose to report on some optional indicators but not all.

The Goals focused on conservation (A) and sustainable use (B) are considerably better covered (90-100% and 67-83% of elements at least partially covered respectively) than those on benefit sharing (C, 0%) and resourcing (D, 20-60%; Figure 3). Additionally, there are gaps in the coverage of the Monitoring Framework for all three sections of the Targets: reducing threats to biodiversity (Targets 1 to 8: 28-47/54, 52-87% at least partially covered); meeting people’s needs through sustainable use and benefit sharing (Targets 9 to 13: 19-24/39, 49-62% at least partially covered); and tools and solutions for implementation and mainstreaming (Targets 14 to 23: 51-56/69, 74-81% at least partially covered; Figure 3).

### 2.2 Headline indicators

It is apparent that headline indicators alone will only allow partial tracking of progress, completely covering 20/190 (11%) and partially covering an additional 29/190 (15%) elements, and relating to only 16/23 Targets. The disaggregations of these indicators, recommended by the AHTEG to enhance their ability to report on elements of the Goals and Targets, would increase coverage by headline indicators to 26/190 elements (14%) completely and an additional 31/190 (16%) elements partially.

The coverage of some headline indicators could be improved by expanding geographic or taxonomic coverage through data acquisition and methodological development (e.g. A.1, A.3 and A.4). Furthermore, several headline indicators (e.g. B.1, 2.1, 3.1 and 6.1) have the potential to cover more elements of their specific Goal or Target if additional disaggregations can be developed (e.g. by ecosystem type or gender). In some cases, the current lack of potential disaggregations results from a methodological challenge (e.g. 12.1), while in others, it is due to data availability (e.g. 6.1). Improving and/or broadening data collection for these indicators would potentially improve the coverage of the Monitoring Framework without the need to develop completely new indicators. Through disaggregation, some indicators may further support management plans targeting specific issues (e.g. 3.1 by areas of importance for biodiversity or by indigenous and traditional territories; A.1 by drivers or by protected areas and OECM), in addition to tracking overall progress towards a Goal or Target.

Some headline indicators have been specifically designed following the adoption of the Monitoring Framework (e.g. B.1, C.1, C.2 and 21.1) and have the potential to cover the respective Goals and Targets comprehensively but are not yet fully in place. Committing the resources needed to finalise, test and implement their methodologies would enable the monitoring of targets that currently cannot be monitored (Figure 3).

Some headline indicators are so narrowly focused that they provide very limited coverage of their target, and component and complementary indicators are needed to track progress effectively. Specifically, Targets 5 and 12 are only sparsely covered due to the narrow focus of headline indicators 5.1 and 12.1 (Figure 3 and Box 2); both are SDG indicators and cannot be amended easily.

Some Targets are likely to require significant effort to fill the gaps identified. Specifically, the headline indicators proposed for Targets 1 and 9 remain hypothetical (Figure 2) and are a high priority for methodological development. In doing so, it may be helpful to identify synergies in data requirements that may simplify indicator compilation. For example, synergies between indicators are possible for Targets 5 and 9 (addressing trade in wild species and benefits derived from the use of wild species, respectively).

### 2.3 Binary indicators

The 14 binary indicators cover 13 Targets, 8 of which have no headline indicator. They will provide a valuable source of information that can be used to rapidly and reasonably assess progress made on taking measures to deliver on Goals and Targets. However, by their very nature, they cannot inform on the realised implementation (e.g. what kind of measures, their breadth or relevance) or outcomes of these measures. As such, they can only track those elements in Goals or Targets that specifically call for actions (e.g. Target 15) and not the elements relating to the results of such actions. Therefore, for all Targets calling for results and for which only binary indicators are available (Targets 8, 14, 16, 20, 22 and 23), only partial coverage of the elements can be achieved with the current Monitoring Framework (Figure 3). Efforts are needed to design fit-for-purpose quantitative indicators that can support the binary indicators to cover the outcome elements of those targets. Collaboration may help lower the burden of designing new indicators. For example, the Women4Biodiversity developed an indicator on the Gender Plan of Action^34^ (Target 23) and the Global Youth Biodiversity Network is working on an indicator that may be suited to report on targets relevant to children and youth (Target 22).

### 2.4 Complete gaps

For 23-56 elements (12-29%, Figure 2), no indicators are available to track progress. There are three Goals (B, C and D) and 11 Targets (1, 2, 3, 4, 10, 11, 12, 13, 17, 18 and 19) containing elements that cannot currently be monitored under the Monitoring Framework (Figure 3). For some of these (e.g. D and 12), the gaps are sufficiently significant that new indicators may need to be added to the Monitoring Framework. For others (e.g. 3 and 13), extending the methods or identifying disaggregation options for existing headline indicators to cover the gaps might be possible. Other Targets (e.g. 2 and 4) may require specific indicators to complement the existing headline indicators. These additional indicators could come in the form of component indicators specifically addressing the gaps identified (e.g. Target 2: “enhancing ecological integrity and connectivity”), or additional straightforward binary indicators may suffice (e.g. Goal D: “adequate capacity-building is secured”). Additional component and complementary indicators could be developed by organisations currently focused on those specific aspects of the Goals and Targets (for example, human-wildlife conflict^35^) whereas additional binary indicators could be proposed by Parties and follow the methodology designed by the AHTEG. Alternatively, Parties may choose to include national indicators addressing these gaps within their national context.

## 3. General recommendations

### 3.1 Improving the coverage of the Monitoring Framework

The Monitoring Framework has some important limitations. However, its adoption is a major step forward compared to previous strategic plans (the previous plan did not have a Monitoring Framework, the use of indicators in national reports was optional and reporting was limited). Moving forward, the Monitoring Framework sets the stage for building and improving national monitoring systems and improving monitoring over time.

These limitations of the Monitoring Framework will be particularly important for countries to consider when they develop their national monitoring systems. Through the adoption of the mechanisms for planning, monitoring, reporting and review, Parties recognized that at the national level, each country should develop a national monitoring plan for monitoring their national biodiversity strategy and action plan (NBSAP) and that this national policy should utilize headline, binary, component, complementary and national indicators as appropriate. Given the significant challenges in developing and implementing the indicators in the Framework, it is highly unlikely that all Parties will utilise all of them, especially as the component and complementary indicators are optional, and the disaggregations of the headline indicators are recommended rather than mandatory. As such, gaps are inevitable and it is important for both Parties and the international community to see the Monitoring Framework as something that can be improved over time through both investments in national monitoring and scientific research. At this time, the biggest risk to effective tracking of global progress is a lack of ambition in implementing the Monitoring Framework.

Our results highlight that relying solely on the indicators required for national reporting will cover only some of the elements in the GBF. The Monitoring Framework could provide good coverage if the recommended disaggregations of the Headline indicators as well as the optional indicators are implemented. For example, the current coverage for Target 3 is entirely provided by disaggregations of headline indicator 3.1. A lack of ambition or resources from Parties could result in few component or complementary indicators or disaggregations being utilised and reported. In such a case, we would have little ability to judge whether most of the Goals and Targets were met in each country and at a global scale: much of the GBF would be unmonitored. Therefore, efforts must be made to ensure that recommended disaggregations and optional indicators are included in national monitoring plans and reported in the national reports.

Particular attention should be paid to all those elements assessed as “potentially” covered. Work is needed to develop appropriate indicators for these 25 elements, but they are not yet ready for national implementation. There is a short-term opportunity to improve coverage of the Monitoring Framework with more limited investment of time and resources. Rapidly completing the development and testing of these indicators would allow Goal C and Targets 6, 11, 13 and 21 to be monitored effectively.

Specific gaps can be addressed at the national level using national indicators that are not included in the current list. Parties may develop these indicators at the national level or international organisations may do so at the global level and then disaggregate to the national level. This approach would not systematically address the gaps in the Monitoring Framework but would improve coverage of the GBF for some Parties and make available additional indicators for consideration in the next rounds of negotiations (it is expected that the list of component and complementary indicators will be updated at COP17 and potentially at future COPs^36^). For example, whilst the monitoring of some aspects of freshwater resources and aquaculture are gaps, some Parties have chosen to report on their status using national indicators^37,38^.

It is unlikely that new headline or binary indicators will be adopted by Parties and added to the Monitoring Framework before the next strategic plan for the Convention is developed in 2030. Further component and complementary indicators, however, may be added to the Monitoring Framework if they meet the criteria for inclusion, but these will remain optional. Therefore, while there are opportunities to fill specific gaps with additional indicators, as discussed above, improving the Monitoring Framework should be seen as a long-term endeavour that will continue over several years. Understanding the underlying reasons for gaps provides an opportunity to improve global biodiversity monitoring efforts. In the following paragraphs, we address the general requirements for an effective Monitoring Framework for today and into the future.

### 3.2 Data needs

Many gaps or issues of partial coverage are linked to data availability. There are historical reasons behind the incomplete coverage of biodiversity data globally^22,39^ which have resulted in poor coverage of many species, genetic diversity and ecosystem extent and integrity around the world, especially in tropical regions^40^. As such, improving sampling efforts of biodiversity data is a key priority for monitoring of the GBF, in particular for indicators A.1, A.4 and 2.1. Data collection, archiving, and reporting efforts should follow the FAIR^41^ and CARE^42^ principles and take advantage of new technology and guidance on designing effective biodiversity monitoring systems to strategically fill gaps and compute indicators^43–46^. They should build on and make use of existing data infrastructures, tools, and community-developed standards that provide well-functioning and evolving mechanisms for sharing and integrating biodiversity data from diverse sources of evidence^47^.

Ideally, biodiversity monitoring systems should be designed to integrate *in situ* observations gathered by conventional research programmes, such as long-term ecological research networks^17,48,49^, or community-based monitoring and information and citizen science efforts^50^ with remote sensing data^51^ and expert judgement, to take advantage of big data^52^ and potential advances in artificial intelligence^53^. To monitor the Goals and Targets of the GBF and link biodiversity outcomes to drivers of change, these data will need to be linked to national census surveys and other relevant physical, economic and social measures. However, gaps exist in these datasets themselves and national census surveys specifically may need to be updated to reflect the needs of the GBF. For example, questions on mainstreaming (Target 14) may help address a data-poor section of the Monitoring Framework. One important reason for this poor and declining^54^ biodiversity data coverage is the lack of investment and institutional support for biodiversity monitoring efforts^55^. However, many of the headline indicators are actively developed and supported by international organisations. If resourced, the same organisations can guide data collection, provide capacity-building, and offer support to Parties required to implement these indicators. Additionally, civil society and volunteer efforts to collect biodiversity data provide significant value to states in administrative cost savings^56^ and should be promoted and integrated with government-led biodiversity monitoring systems.

Collecting additional relevant data could further allow disaggregation of indicators, improving the Monitoring Framework’s coverage. However, it should be noted that there is a difference between asking Parties for more disaggregated data (which would increase the burden of reporting) and overlay of data provided by Parties with information from other sources (which implies more effort in analysing collated data). Ultimately, the issue of data collection, whether done by Parties or in collaboration with non-governmental organisations, is one of capacity^57^. Resourcing additional data collection efforts will be a significant challenge for many Parties and will require North-South, South-South and triangular collaboration as well as financial resources to be made available.

### 3.3 Methodological needs

The issue of data is linked to the tools and methods currently available to detect trends in biodiversity and benefits derived^58^. Adapting the way biodiversity change is understood to better reflect the data available today is necessary to avoid difficulties in measuring change^59^. Furthermore, the scientific community has not yet come to a consensus on understanding and monitoring different facets of biodiversity and people’s needs^24,60^. Indeed, the needs of the GBF go beyond what might have historically been considered biodiversity data and require that information on drivers of biodiversity change and societal outcomes be monitored (e.g. for Targets 9, 11 and 12), potentially requiring cross-sectoral data-sharing agreements. In some cases, this results in the need for entirely novel data collection and analytical methods, such as for financial flows^61^ (indicators D.1, D.2 and D.3) and access and benefit sharing (ABS)^62^ (indicators C.1 and C.2). Specifically for ABS indicators, legal frameworks (e.g. the Nagoya Protocol^63^) did not foresee the need to measure societal outcomes but rather national compliance metrics. Additionally, social processes related to participation, equity and rights (e.g. for Targets 22 and 23) also need effective monitoring to ensure that all parts of the GBF are effectively implemented. Thus, the Monitoring Framework now requires new types of biodiversity-related data and analytical methods to enable policymakers to evaluate whether and to what extent the third objective of the CBD is being met.

Integrating across data sources and between social and ecological data is a significant challenge that requires well-designed pipelines and data products that can standardise information and make it usable for indicator calculation. For example, essential variables for biodiversity^64,65^ and ecosystem services^66^ can standardise multiple data types, enabling the use of analytical pipelines to compile indicators and support further development efforts. Additionally, approaches to recognise and support the role of Indigenous and local knowledge as complementary to the convential scientific processes are improving^67,68^ but more remains to be done for these to be systematically applied and fully inclusive^69^.

One of the strengths of the Monitoring Framework is that it will enable consistent reporting of indicators by countries, supporting comparisons across countries and aggregation for regional and global synthesis. For example, the AHTEG recommended a consistent approach to compiling and reporting indicators related to ecosystems, their protection, restoration, sustainable use, and status. This would allow the impacts of the actions (e.g. restoration indicator 2.1) to be tracked on outcomes, such as extent of natural ecosystems (indicator A.2) and their risk status (indicators A.1 Red List of Ecosystems). The AHTEG recommended using the IUCN Global Ecosystem Typology (IUCN GET)^70^, endorsed in March 2024 as an international statistical classification by the UN Statistical Commission^71^, to support consistent reporting and disaggregation of ecosystem-related indicators by ecosystem functional group (level 3 in the typology). To do so, countries would cross-reference their national ecosystem classifications and maps with the IUCN GET to enable reporting based on national ecosystem data and assessments. Significant progress has been made by Parties to map ecosystems to date^72^ and tools are available to facilitate advancement^73^. Accelerating its adoption would potentially allow a complete picture of given ecosystem groups (e.g. coral reefs, or tropical lowland rainforests) within a country, globally and across the indicators.

An important focus area for rapid and systematic indicator calculation is interoperability^74,75^. Designing analytical pipelines capable of fully integrating and interpreting data types and meaning^76,77^ and producing indicator calculations will significantly reduce the capacity needs of Parties^78^. Prioritising these efforts to compile headline indicators and share code openly with all Parties could significantly reduce the equity and capacity barriers to monitoring the GBF and free up resources for Parties to include additional optional indicators relevant to their needs. Some of the infrastructure to enable this process is already in place^79–81^ but additional efforts are required to fully operationalise it.

Finally, accounting for, handling of, and reporting on uncertainty in biodiversity estimates and indicators will be essential to effectively guide decision-making and implementation of the GBF. Uncertainty in trend detection capabilities has shed doubt on understanding global biodiversity change^82^ and on the ability to detect improvement or further decline in biodiversity with a high level of confidence^59^. One solution is to use modelling techniques, such as Bayesian inference^59^, that can allow updated estimates of progress as new data and evidence arise within a robust and rigorous framework for dealing with and reporting on uncertainty^59,83^. Communicating uncertainty in indicator values to decision and policy makers is crucial for the Monitoring Framework to go beyond progress tracking and effectively support implementation.

### 3.4 Building the infrastructure to monitor biodiversity globally

Long-term monitoring to fill data gaps, streamlining indicator compilation, capacity-building and integrating knowledge systems is a global challenge. The GBF is a common endeavour, that, although Party-led, requires nations and stakeholders within them to work together^84^. The upcoming fast-track assessment by IPBES on monitoring biodiversity and nature’s contributions to people will assess current capacity, identify needs and suggest solutions^85^. One solution is to support international cooperation by establishing a Global Biodiversity Observing System (GBiOS). A GBiOS would support the implementation of national biodiversity monitoring networks^43,46,86,87^ while assembling an international system similar to that used to monitor the oceans (i.e. the Global Ocean Observing System) or trends in climate and weather (the WMO Integrated Global Observing System). These systems allow countries to opt in and thus benefit from funding and shared resources and technologies, such as data and modelling infrastructure^88^. Following this model, GBiOS could therefore exist as a federated international network of existing and new national monitoring systems^75,87^. Over time, GBiOS could allow multilevel assessments of trends in different facets of biodiversity (genetic, species and ecosystem) that could contribute to calculating indicators.

Headline indicator 21.1, on biodiversity information for monitoring the GBF is currently under development (Figure 2). At the highest level, this indicator could be evaluated if countries reported the number of headline indicators where national datasets, models, monitoring schemes and information tools are available and used. If aggregated and calculated over time this would capture within and across country-level trends in the access and use of data for governance, management and communication of biodiversity outcomes identified by NBSAPs. Development of Indicator 21.1 along three dimensions would measure the coverage in space and time of used and accessible data capturing trends in different biodiversity dimensions, the recognition and use of Indigenous and local knowledge of biodiversity, and the quantity and scope of active biodiversity monitoring activities and the data they produce to support the ready calculation of all relevant headline indicators.

This effort could be supported long-term by information made available by GBiOS, by filling gaps in data coverage that hinder easy calculation of headline 21.1 (Figure 3). Other indicators (e.g. A.2, B.1, 2.1, 3.1) could rely on data collected and shared through GBiOS, addressing gaps in geographical coverage and data availability. Making GBiOS a reality requires (i) a sustainable and inclusive governance model allowing countries to opt in and actively participate, (ii) a funding mechanism to support Parties looking to invest in their national biodiversity observation networks, and (iii) the human capacity and technical infrastructure required for GBiOS to connect countries into an operational international network^33^. When brought together with other knowledge types, including Indigenous and traditional knowledge, scientific expert judgement, socio-economic data, and information on governance and regulation, a GBiOS could generate the information needed to guide investments in monitoring capacity to fill the gaps in the Monitoring Framework.

Such a system would further promote synergies between the CBD and the global climate agenda^89^. Climate monitoring systems are more advanced than those of biodiversity yet both crises are closely linked^90,91^. Some important differences between climate and biodiversity have limited efforts to connect these Conventions and their data collection systems: biodiversity metrics are less standardised^64^, their scale of change is more local and sensitive to small-scale changes^92^, methods for data collection are more diverse^45^, and the global policy relevance of the biodiversity crisis has lagged behind that of the climate crisis^93,94^. Yet, there are many synergies between monitoring climate and biodiversity (e.g. nature-based solutions are an important component of climate policy^95,96^); connecting them would improve understanding of biodiversity change and support efficient data collection efforts^87^. Taking advantage of the capacity and knowledge already built in the climate space may also further the implementation of national and global biodiversity observing systems. Other sectors, such as those carrying out environmental impact assessments, also produce important knowledge and could contribute to monitoring of outcomes under the GBF^97^.

### 3.5 Cross-cutting considerations

Section C of the GBF outlines the cross-cutting considerations for the GBF, including its implementation and evaluation. It requires that the implementation of the GBF be consistent, inter alia, with the rights and contributions of IPLCs, different value systems, human rights, gender, intergenerational equity, and human health^7^. The considerations in Section C are partly, but not wholly, addressed by the existing Monitoring Framework and its headline and binary indicators. No indicators were specifically designed nor included to measure progress towards Section C. However, the current headline and binary indicators can be used as informative proxies. Indeed, some indicators are particularly relevant (e.g. 9.1, 14.b, 22.b and 23.b), while others invite that additional data be collected to allow relevant disaggregation (e.g. for IPLCs/women/youth for A.1, B.1, C.1 and C.2). In many cases, the necessary disaggregation of headline indicators is not yet possible to show how the cross-cutting aspects of the GBF are being delivered due to a combination of limitations in data and methodological approaches.

The considerations can be partly addressed by using indicators relevant for the defined social groups in Section C. Specifically for IPLCs, the four traditional knowledge indicators^98^ could be used to complement the headline and binary indicators. Within the Monitoring Framework, the traditional knowledge indicators are component or complementary indicators for certain targets and can provide a basis for disaggregating existing headline indicators (e.g. for indicator 3.1 by disaggregating spatial headline indicators by Indigenous and traditional territories).

A human rights-based approach to monitoring has well-established guidance, methodology, and indicators^99,100^. This approach is applied to monitoring of the traditional knowledge indicators and could be extended to the whole Monitoring Framework. Additionally, to meet existing commitments, particular attention is needed to address disaggregation by gender, age, disability, IPLCs, education, and socio-economic status. Although data availability may limit the use of disaggregated data in the next national reports, the gaps, priorities, and opportunities for data disaggregation and participatory approaches should be identified and advanced to improve future monitoring and resulting management action.

Monitoring can also be adapted to include participatory processes, including through involving a broad range of sources in acquiring the underlying data needed to populate the indicators. The Monitoring Framework invites Parties and relevant organisations to support community-based monitoring and information systems (CBMIS). The contribution of CBMIS to monitoring progress and achieving biodiversity targets is now well documented^101^. Some community-based monitoring tools implementing a human rights-based approach, such as the Indigenous Navigator^102^, are already operational. CBMIS provide cross-cutting data and knowledge useful for monitoring multiple Targets and Section C^103^. Specifically, CBMIS collect data in remote areas where gaps are common even for well-developed indicators. Additionally, CBMIS often provide traditional knowledge that can be invaluable in promoting conservation outcomes and the implementation of the GBF^68,104^. Furthermore, CBMIS are typically community-focused and reflect the needs and priorities of their members as well as different value systems^105^. Typically, CBMIS also collect additional information on health and human rights outcomes. As such, these monitoring systems should be an integral part of data collection efforts wherever they exist. Finally, community-based monitoring efforts in collaboration with scientists have proven to be a cost-effective way of furthering understanding and managing biodiversity^106,107^ and therefore should be supported by all Parties.

Nevertheless, there is a need for more work to be done across all the cross-cutting elements of Section C as gaps remain in our understanding of how these topics interact with biodiversity. This was not assessed in detail in the gap analysis and remains an open area for research. This work is essential given that Parties often noted difficulties in taking Section C into account in the development of their national targets and only around one-third of the national targets submitted by Parties by October 2024 contained information on how the issues identified in Section C were being taken into account^108^.

## 4. Next steps in implementing the Monitoring Framework

### 4.1 Reviewing and updating the Monitoring Framework

The work of the AHTEG, in collaboration with many organisations and individuals, has significantly improved the readiness level of existing indicators and the ability of Parties to use the Monitoring Framework to track progress towards the ambitions of the GBF^8^. Work on improving those indicators currently in the Monitoring Framework should be encouraged and appropriate organisations resourced. Following the next round of reporting, there will be an opportunity at COP17 to review how the Monitoring Framework has been operationalised at the national level by reviewing what was reported by Parties and considering whether it needs further development. This engagement is key to the success of the Monitoring Framework.

The Monitoring Framework is expected to be kept under review, providing an opportunity for updates and improvements. A formal process of update and review for the Monitoring Framework, akin to the annual refinement and quinquennial comprehensive review of the Sustainable Development Goals indicators^26^, but focusing on updating the list of indicators as well as their methodology, could be established to fill gaps and improve the Monitoring Framework. Such a process could learn from the AHTEG and include experts and stakeholders representative of the GBF with a clear mandate and resources to update the Monitoring Framework.

### 4.2 Addressing resourcing and capacity-building priorities

The most likely barrier to a successful implementation of the Monitoring Framework is one of human and financial capacity. Issues of equity are particularly relevant in this context as areas with the most biodiversity are often those with the least capacity to monitor its status^109^. Without the resources to implement the required indicators in the Monitoring Framework, many Parties will not be able to report on them and gaps will remain unfilled — thus, uncertainties about progress under the GBF will persist. Through COP decisions, Parties have committed to prioritising biodiversity planning and monitoring nationally. Financial resources are now needed to establish and strengthen national and regional monitoring frameworks. These resources should be used to build nationally relevant biodiversity observation systems with the people, technologies, and knowledge systems needed to implement the Monitoring Framework.

The challenge in funding monitoring will be particularly acute for those Parties with the least resources, for which multilateral and innovative funding sources may be required. Private and public cooperation, through blended finance for example^110^, can potentially help mobilise some of these funds. While governments will need to invest more into monitoring, private sector actors also have a role in enhancing funding levels and deploying innovative solutions. Around half of the world’s GDP is moderately to highly dependent on biodiversity^111^, meaning that private companies and financial institutions have a considerable stake in this issue. Direct engagement of the private sector in supporting biodiversity monitoring efforts^112^ must respect the rights of all communities involved and ensure that all data are collected fairly and equitably. Additionally, the implementation of benefit-sharing agreements under the CBD and its Nagoya Protocol, as well as the newly-created multilateral mechanism for the fair and equitable sharing of benefits from the use of digital sequence information on genetic resources^113^, may help build capacity among those Parties where the genetic resources are being sourced^114^, as well as among IPLCs. However, some experts have argued the opposite may occur if ABS legislation is overly burdensome or bureaucratic^115^.

Finally, beyond national efforts, global cooperation is required to resource the organisations that are supporting methodological research and capacity building around the indicators. This is essential for the full development of indicators and their disaggregation, for enhancing harmonization among indicator frameworks when relevant (e.g. UNFCCC work on indicators under the Global Goal on climate adaptation), and for increasing the coverage of indicators that support the effective implementation of the Monitoring Framework over the long-term.

## 5. Conclusion

The adoption of the Monitoring Framework was a monumental achievement. This decision by Parties represents a shift in how transparency and responsibility will be assured. The Convention has moved away from national reports containing largely qualitative narrative, to reports requiring quantitative indicators. Despite its existing limitations, the Monitoring Framework marks a paradigm shift in the implementing the CBD. Further developing and implementing it will be challenging, but changing the status quo is never easy.

The Monitoring Framework has been developed considerably since its first draft was negotiated^7^. The present analysis shows that, despite the gaps identified, there is a large quantity of data, indicators and knowledge is available to implement the Monitoring Framework, making use of the guidance and resources available. It now stands ready to inform on progress towards achieving global Goals and Targets in a way that has not been previously possible in the CBD’s history. However, its effectiveness will depend on the degree to which Parties implement it. While all Parties must balance multiple priorities and face practical, technical, financial and social constraints on what can realistically be monitored, the ambition of Parties in implementing monitoring and reporting indicators will determine our ability to judge progress towards the Goals and Targets, and, ultimately, whether they have been achieved. The Monitoring Framework provides an opportunity to develop and invest in national biodiversity monitoring systems and to improve national and global data sharing and connection of monitoring systems.

The academic and NGO communities have roles to play in supporting implementation by providing capacity building, training and data-informed workflows, which are required to lower the barriers to the implementation of the Monitoring Framework, especially for developing nations. Academic and citizen scientists, NGOs and IPLCs have an important role to play in collecting data and compiling indicators to help Parties report on the many indicators of the Monitoring Framework. Whilst the gaps identified here are concerning and should be addressed, a failure to implement the Monitoring Framework will result in a much greater number of gaps. We strongly encourage the scientific community and other relevant actors to engage with and support Parties in their efforts to implement the Monitoring Framework. Its implementation through the 7th national reports expected in 2026 will provide a first opportunity to gauge the adequacy of monitoring and the extent of progress towards achievement of the GBF.

The Monitoring Framework will be essential to delivering on the ambition of the GBF by 2050. It should be considered a global learning mechanism, allowing enough flexibility for Parties to meet their objectives while sharing progress and experiences globally to improve outcomes for the whole of society. The achievement of the GBF Targets for 2030 and the Vision and Goals for 2050 depends not only on the comprehensive and effective implementation of policies and actions but also on effective implementation of the Monitoring Framework.

## 6. Methods

### 6.1 The work of the AHTEG

#### 6.1.1 Headline indicators

Methodologies for compiling headline indicators were reviewed, revised, or, in some cases, developed *de novo* by the AHTEG, in consultation with relevant organisations and experts. Metadata covering the rationale, methods, available data, and possible disaggregations for compiling each indicator are available for all headline and binary indicators agreed at COP15 except for two (Figure 2)^116^.

The work of the AHTEG started with the evaluation of the metadata for each indicator by sub-teams of experts on each topic. The existing metadata for each indicator was assessed for completeness and whether it clearly reflected the latest guidance and methodology for indicator calculation. This also involved assessing whether the data required to calculate the indicator were freely available for all countries. Shortfalls in methodology, data, and the description of the indicator were considered and documented for each indicator. Depending on the levels of development of the indicators, AHTEG members worked directly with agencies supporting methodological research and capacity building for the indicators to revise the meta-data, providing advice, text edits and methodological revisions where needed, and in some cases, the development and initial testing of new indicators.

#### 6.1.2 Binary indicators

The selection of questions and evaluation of potential answers involves careful consideration if binary indicators are to provide meaningful metrics of progress towards the Goals and Targets. The AHTEG reviewed the proposed text of binary indicator questions and answers to produce Goal and Target specific guidance on interpreting questions and answers. This guidance is outlined in the metadata of each binary indicator^116^. Additionally, the AHTEG developed a methodology to aggregate responses across questions using mutually exclusive answer combinations to produce an overall score for each Party that can be used to track global progress towards implementing the measures required to deliver on the GBF.

#### 6.1.3 Qualitative cross-cutting considerations

Section C of the GBF outlines a set of “cross-cutting” considerations to address in the implementation of the GBF and which are relevant to the Monitoring Framework. Some of these cross-cutting considerations (namely human rights, gender, IPLCs, and the ecosystem approach) were reviewed by the AHTEG within the context of the Monitoring Framework and its indicators. Specific guidance^116^ was provided for considering human rights and ecosystem approaches and how these could be implemented within the Monitoring Framework. Current limitations in the ability of the Monitoring Framework to track progress towards the cross-cutting considerations were also assessed.

### 6.2 Gap analysis of the Monitoring Framework

#### 6.2.1 Gap analysis through expert elicitation

Gap analyses are used to identify areas where knowledge, data or action are lacking relative to a desired outcome or standard^117–120^, in this case the ability to monitor comprehensively the GBF’s Goals and Targets. Gap analyses are common practice in fields where action needs to be taken, resources are limited and knowledge is incomplete, as is often the case in conservation^29,30,121^ and resource management^122,123^. A common method to carry out a gap analysis is through the use of expert elicitation^123–125^.

Experts, through formal training, acquired skills and experience, are deemed more able to critically assess a situation and identify where gaps exist^126^. We refer to experts as “people who are considered by their peers and society at large to have specialist knowledge and who are consulted to make an estimate”^32^. However, expert judgements are dependent on the kinds of experts convened and the way their knowledge is elicited^32^.

The process of selection of the AHTEG is described in the terms of reference for the AHTEG, specifically ensuring representation of different areas of technical expertise and ensuring balance in expertise on all aspects of the goals and targets of the Kunming-Montreal Global Biodiversity Framework, also taking into account geographical representation, and the representation of indigenous peoples and local communities, women’s and youth groups, and major stakeholders, gender balance and the special conditions of developing countries, in particular the least developed countries and small island developing States, and countries with economies in transition, also taking into consideration the special situation of developing countries that are most environmentally vulnerable, such as those with arid and semi-arid zones, coastal and mountainous area^27^.

The potential limitations of expert judgement can be further mitigated through the use of well-established structured procedures^126^. Here we followed a multi-step process using individual reflection, focus groups and broad discussion over the course of several months, detailed below.

#### 6.2.2 Mapping out the ambition

The AHTEG was mandated to analyse whether the indicators in the Monitoring Framework could track progress towards the GBF’s Goals and Targets. This task required analysis of the full ambition of the GBF as reflected in the wording of the Goals and Targets agreed by governments following the lengthy negotiations through the CBD. The text of each Goal and Target is often long and complex, containing multiple clauses and components that are each important to deliver on the ambition of the GBF. To enable a comprehensive and holistic analysis of the Monitoring Framework’s ability to track progress towards the whole GBF, the text of each Goal and Target was first divided into a set of distinct elements. This built upon work that the co-chairs of the Open-Ended Working Group on the post-2020 Framework conducted on identifying elements of each goal and target. Elements were defined as those components of the Goal or Target that would need to be measured and reported independently to evaluate whether the Goal or Target had been fully achieved. The rationale for this approach built on the methodology used to assess progress made towards the Aichi Biodiversity Targets, which concluded that none of the targets had been fully met, but some had been partially met based on achievement of one or more elements within those targets^5^. The elements were defined exclusively by the text of the Goal or Target, without the relevant considerations from Section C of the GBF.

This exercise was done in two steps: an initial list of elements across most of the Goals and Targets was proposed by one AHTEG member and was shared online with all other AHTEG members for review and commentary. All AHTEG members were then invited to make suggestions to further split, combine or redefine the elements, based on their specific knowledge of different aspects of the GBF. A consolidated list of elements was thus agreed by all AHTEG members (Supplementary Table 2). Separately, three expert groups, also involved in the development of specific indicators in the Monitoring Framework, were invited to propose elements relating to the remaining Goals and Targets on which they were focussed: specifically, the Liaison Group on the Cartagena Protocol for Target 17, the Informal Advisory Group on Technical and Scientific Cooperation for Target 20, and the Technical Expert Group on Financial Reporting for Goal D and Targets 18 and 19.

Due to the complexity of the language of some Goals and Targets, incorporating multiple commitments including both actions and outcomes, the number of elements for each varies considerably. For example, Target 12 contains 8 such elements (Box 2; see Supplementary Table 2 for the complete list of elements for all Goals and Targets). The number of elements ranged from 4 for Target 23 to 14 for Target 22, with a median of 6 and a total of 190 elements across all Goals and Targets (Supplementary Table 3).

#### 6.2.3 Assessing coverage

The indicators in the Monitoring Framework were then mapped to the elements of each Goal and Target to identify how comprehensively the GBF could be monitored, which elements were not covered by the headline indicators, their disaggregations, binary indicators, and component or complementary indicators (Box 2). Note that the cross-cutting considerations set out in Section C of the GBF were not considered in the coverage of the indicators. The results and explanations for the coverage status of each element were made available to Parties ahead of SBSTTA26^33^.

This process was conducted through a two-step direct elicitation process^29,127^: first, a remote elicitation^31,128^ step via a survey shared with all AHTEG members prior to the AHTEG’s 6^th^ meeting in Cambridge (March 2024) and second, a structured facilitated working session^129^ at that meeting.

The survey’s focus was broadly to identify where the headline and binary indicators covered the elements of the Goals and Targets and where component and complementary indicators might support monitoring of the elements. Respondents were asked to select those elements that they thought could be monitored using each headline and binary indicator in turn, then they were given the option to repeat this for any complementary or component indicator of their choice. At this stage, respondents did not have access to the updated metadata and list of component and complementary indicators. As such, respondents were specifying whether the intent and concepts behind the indicators could map onto each element rather than realised indicators with methodology. Therefore, the survey results were not used to report on the gaps in the Monitoring Framework but rather to guide the in-person session in Cambridge.

At the meeting in Cambridge, after all the indicators and their methodologies had been reviewed, the AHTEG members discussed the results of the remote elicitation step during an inception meeting^31^. The results of the survey were presented by a facilitator through a series of graphs. The facilitator then proceeded to explain how the in-person process would be split between a focus group session, where AHTEG members would be given a set of questions to work on in small groups, followed by a larger group session to review the results of the focus groups and discuss any divergence in opinions between experts^31^.

During the focus group session, AHTEG members were asked to split up into self-directed groups of expertise, focusing on specific goals and targets. Within each group, AHTEG members were given the same task with clear questions to review all the Goal or Target elements individually. For each element, groups were asked to list the headline or binary indicator (including recommended disaggregations of headline indicators where relevant) that would allow tracking of progress on the element. When a headline or binary indicator was assigned to an element, groups were asked to qualify the coverage of each element by the corresponding indicator with one of three options:

- “Covered”: i.e. an indicator has been developed and tested in at least some countries and can inform progress on the element
- “Partially”: i.e. an indicator has been developed and tested in at least some countries and can inform progress on part of the element, without fully capturing the full scope of the element being addressed (i.e. it indirectly addresses the element – tracking legislation put in place but not its impact – or it only addresses the element in part – tracking harvest of fish species rather than wild species)
- “Potentially”: i.e. an indicator for the element is still under development or has not been tested but once completed and tested, is expected to be able to track progress on the element (i.e. headline indicators with methodology in development (Figure 2) or developed indicators with missing disaggregations that could inform on the element but are not currently available)

For each of these categorisations, participants were asked to provide a brief explanation (justification and caveats) for their choice and specify, in the case of headline indicators, whether a specific disaggregation is required to inform progress on the element. For elements where no headline nor binary indicator could inform progress, participants were asked to suggest component or complementary indicators that may be appropriate to inform on progress towards these elements.

This process was repeated after the meeting in Cambridge for specific indicators that had not received sufficient attention during the meeting. These subsequent exercises were conducted with AHTEG members for targets 14 and 16, with the Technical Expert Group on financial indicators for goal D and targets 18 and 19 and with relevant members of the Secretariat for targets 17 and 20.

The results of these processes were analysed and compiled into a report^116^. These were shared with all participants prior to submission for review and feedback. Participants were asked to review the results individually once more to confirm these accurately represent the consensus opinions of the AHTEG.

Following SBSTTA’s 26^th^ meeting in Nairobi (May 2024), the results of the gap analysis were updated to reflect the new language of the binary indicators^130^.

#### 6.2.4 Limitations

It is important to acknowledge that although the gap analysis was conducted by experts under a formal structured process, many of whom were actively involved in the design of the methodology for indicators and the overall Monitoring Framework, the results reflect their judgements at the time the exercises were carried out. Repeating this process with different individuals having access to the same information may yield somewhat different results. However, few, if any, people were more well-versed in the Monitoring Framework than the members of the AHTEG when this work was conducted, and their judgement was recognised and used as the basis for decision-making and recommendations at SBSTTA 26^132^ and COP16^131^. As such, this is the best information available about where the Monitoring Framework stands and what is needed going forward.

#### 6.2.5 Updates to the Monitoring Framework following COP16

While the meeting of COP16 in Cali, Colombia, did not finalise an agreement on the updates to the Monitoring Framework, negotiations on the indicators resulted in the possible editing of one headline indicator, the addition of one headline and one binary indicator and the removal of four component indicators^131^. Indicator 7.2 was singled out for a change that remains to be decided. The new headline indicator 22.1 tracks land-use change and land tenure in the traditional territories of indigenous peoples and local communities. The new binary indicator 5.b asks Parties about the existence of legal instruments or other policy frameworks to regulate trade in wild species. Neither of these indicators was present in the Monitoring Framework when the gap analysis was carried out^132^. Therefore, the potential increase in coverage from their addition is not included in this work. Additionally, all four component indicators that were removed were for Target 18: revenue generated from biodiversity-relevant taxes, fees and charges; monetary value of biodiversity-positive subsidies; number of other positive incentives in place for biodiversity (by type); monetary value of other positive incentives in place for biodiversity. Removing these component indicators will likely reduce the coverage of the Monitoring Framework for Target 18, which is not reflected in our results.

## 8. Acknowledgements

We thank all members of the AHTEG and other experts who supported the AHTEG in its work for their time, input, and commitment to the process. We extend our thanks to the Secretariat of the Convention on Biological Diversity for their help in organising the AHTEG’s meetings and producing the supporting documents for SBSTTA and COP. Further, we thank UNEP-WCMC for their support in facilitating the AHTEG’s work, especially during its meeting in Cambridge. We also thank the governments of the United Kingdom and Canada as well as the European Union for their funding in support of the work of the AHTEG. A.G. acknowledges the support of the Liber Ero Chair in Biodiversity Conservation. T.S. acknowledges the support of The Pew Charitable Trusts for travel to the AHTEG meeting in Cambridge.

## 9. Data availability statement

Summary data used to produce the figures and results in this article are publicly available here: https://figshare.com/s/ae2e94d993e078b0aaba

## 10. Disclaimer

The views expressed in this manuscript are those of the authors and do not necessarily reflect the views of the organizations by which those authors are employed.

## References

1. United Nations. Convention on biological diversity. (1992).

2. CBD. COP Decisions. https://www.cbd.int/decisions (2024).

3. Butchart, S. H. M., et al. Chapter 3. Assessing Progress towards Meeting Major International Objectives Related to Nature and Nature’s Contributions to People. https://zenodo.org/doi/10.5281/zenodo.3832052 (2019) doi:10.5281/ZENODO.3832052.

4. Buchanan, G. M., Butchart, S. H. M., Chandler, G. & Gregory, R. D. Assessment of national-level progress towards elements of the Aichi Biodiversity Targets. Ecological Indicators 116, 106497 (2020).

5. Secretariat of the Convention on Biological Diversity. Global Biodiversity Outlook 5. (2020).

6. CBD. Kunming-Montreal Global biodiversity framework. (2022).

7. CBD. Monitoring framework for the Kunming-Montreal global biodiversity framework. (2022).

8. Affinito, F., Williams, J. M., Campbell, J. E., Londono, M. C. & Gonzalez, A. Progress in developing and operationalizing the Monitoring Framework of the Global Biodiversity Framework. Nat Ecol Evol (2024) doi:10.1038/s41559-024-02566-7.

9. Tomoi, H., Ohsawa, T., Quevedo, J. M. D. & Kohsaka, R. Is “Common But Differentiated Responsibilities” principle applicable in biodiversity? – Towards approaches for shared responsibilities based on updated capabilities and data. Ecological Indicators 145, 109628 (2022).

10. Watson, J. E. M. et al. Talk is cheap: Nations must act now to achieve long-term ambitions for biodiversity. One Earth 4, 897–900 (2021).

11. Maney, C. et al. National commitments to Aichi Targets and their implications for monitoring the Kunming-Montreal Global Biodiversity Framework. npj biodivers 3, 6 (2024).

12. Butchart, S. H. M., Di Marco, M. & Watson, J. E. M. Formulating Smart Commitments on Biodiversity: Lessons from the Aichi Targets. Conservation Letters 9, 457–468 (2016).

13. Hoban, S. et al. How can biodiversity strategy and action plans incorporate genetic diversity concerns, plans, policies, capacity, and commitments? Preprint at 10.32942/X2PG79 (2024).

14. Parr, T. W., Sier, A. R. J., Battarbee, R. W., Mackay, A. & Burgess, J. Detecting environmental change: science and society—perspectives on long-term research and monitoring in the 21st century. Science of The Total Environment 310, 1–8 (2003).

15. Lindenmayer, D. B. & Likens, G. E. Adaptive monitoring: a new paradigm for long-term research and monitoring. Trends in Ecology & Evolution 24, 482–486 (2009).

16. Lindenmayer, D. B. & Likens, G. E. The science and application of ecological monitoring. Biological Conservation 143, 1317–1328 (2010).

17. Lindenmayer, D. B. & Likens, G. E. Effective Ecological Monitoring. (CSIRO Publishing, 2018).

18. Faith, D., et al. Bridging the biodiversity data gaps: Recommendations to meet users’ data needs. Biodiversity Informatics 8, (2013).

19. Meyer, C., Weigelt, P. & Kreft, H. Multidimensional biases, gaps and uncertainties in global plant occurrence information. Ecology Letters 19, 992–1006 (2016).

20. Etard, A., Morrill, S. & Newbold, T. Global gaps in trait data for terrestrial vertebrates. Global Ecology and Biogeography 29, 2143–2158 (2020).

21. Gallagher, R. V. et al. Global shortfalls in threat assessments for endemic flora by country. PLANTS, PEOPLE, PLANET 5, 885–898 (2023).

22. Chapman, M. et al. Biodiversity monitoring for a just planetary future. Science 383, 34–36 (2024).

23. Nicholson, E. et al. Roles of the Red List of Ecosystems in the Kunming-Montreal Global Biodiversity Framework. Nat Ecol Evol 8, 614–621 (2024).

24. Bennett, E. M. et al. Linking biodiversity, ecosystem services, and human well-being: three challenges for designing research for sustainability. Current Opinion in Environmental Sustainability 14, 76–85 (2015).

25. CBD. Recommendation Adopted By The Subsidiary Body On Scientific, Technical And Technological Advice. (2019).

26. United Nations. Resolution adopted by the General Assembly on 6 July 2017. (2017).

27. CBD. Selected experts for the Ad hoc Technical Expert Group on Indicators for the Kunming-Montreal Global Biodiversity Framework. (2023).

28. CBD. Recommendation adopted by the Subsidiary Body on Scientific, Technical and Technological Advice on 19 October 2023. (2023).

29. Martin, T. G. et al. Eliciting Expert Knowledge in Conservation Science. Conservation Biology 26, 29–38 (2012).

30. Wildermann, N. et al. Informing research priorities for immature sea turtles through expert elicitation. Endang. Species. Res. 37, 55–76 (2018).

31. Hemming, V., Burgman, M. A., Hanea, A. M., McBride, M. F. & Wintle, B. C. A practical guide to structured expert elicitation using the IDEA protocol. Methods Ecol Evol 9, 169–180 (2018).

32. Burgman, M. A. Trusting Judgements: How to Get the Best out of Experts. (Cambridge University Press, 2015). doi:10.1017/CBO9781316282472.

33. CBD. Guidance on needs related to the implementing the monitoring framework of the Kunming-Montreal Global Biodiversity Framework. (2024).

34. Factsheet - Indicators for the Post 2020 Global Biodiversity Framework. Indicator Repository https://gbf-indicators.org/metadata/other/23-1-C (2024).

35. IUCN SSC HWCCSG. Information Document and Discussion Summary regarding the Indicator for Human-Wildlife Conflict in Target 4. (2022).

36. CBD. Mechanisms for planning, monitoring, reporting and review, including the global review of collective progress in the implementation of the Kunming-Montreal Global Biodiversity Framework to be conducted at the seventeenth and nineteenth meetings of the Conference of the Parties. (2024).

37. National Parks & Wildlife Service. Ireland’s 4th National BIodiversity Actional Plan 2023-2030. (2024).

38. Gouvernement Français. Stratégie Nationale Biodiversité 2030. (2024).

39. Boakes, E. H. et al. Distorted Views of Biodiversity: Spatial and Temporal Bias in Species Occurrence Data. PLoS Biol 8, e1000385 (2010).

40. Hughes, A. C. et al. Sampling biases shape our view of the natural world. Ecography 44, 1259–1269 (2021).

41. Wilkinson, M. D. et al. The FAIR Guiding Principles for scientific data management and stewardship. Sci Data 3, 160018 (2016).

42. Carroll, S. R. et al. The CARE Principles for Indigenous Data Governance. Data Science Journal 19, 43 (2020).

43. Pereira, H. & Cooper, D. Towards the global monitoring of biodiversity change. Trends in Ecology & Evolution 21, 123–129 (2006).

44. Pereira, H. M. et al. Global biodiversity monitoring. Frontiers in Ecology and the Environment 8, 459–460 (2010).

45. Scholes, R. J. et al. Building a global observing system for biodiversity. Current Opinion in Environmental Sustainability 4, 139–146 (2012).

46. GEO BON. GEO BON Strategic Plan 2022-2026. (2022).

47. CODATA: Committee on Data of the International Science Council, Uhlir, P. & Pfeiffenberger, H. Twenty-year review of GBIF. (2020) doi:10.35035/CTZM-HZ97.

48. Vaughan, H., Brydges, T., Fenech, A. & Lumb, A. Monitoring Long-Term Ecological Changes Through the Ecological Monitoring and Assessment Network: Science-Based and Policy Relevant. 26 (2001).

49. Ellingsen, K. E., Yoccoz, N. G., Tveraa, T., Hewitt, J. E. & Thrush, S. F. Long-term environmental monitoring for assessment of change: measurement inconsistencies over time and potential solutions. Environ Monit Assess 189, 595 (2017).

50. Chandler, M. et al. Contribution of citizen science towards international biodiversity monitoring. Biological Conservation 213, 280–294 (2017).

51. Cord, A. F. et al. Priorities to Advance Monitoring of Ecosystem Services Using Earth Observation. Trends in Ecology & Evolution 32, 416–428 (2017).

52. Hampton, S. E. et al. Big data and the future of ecology. Frontiers in Ecology and the Environment 11, 156–162 (2013).

53. Silvestro, D., Goria, S., Sterner, T. & Antonelli, A. Improving biodiversity protection through artificial intelligence. Nat Sustain 5, 415–424 (2022).

54. Oliver, R. Y., Meyer, C., Ranipeta, A., Winner, K. & Jetz, W. Global and national trends in documenting and monitoring species distributions. Preprint at 10.1101/2020.11.03.367011 (2020).

55. Moussy, C. et al. A quantitative global review of species population monitoring. Conservation Biology 36, e13721 (2022).

56. Levrel, H. et al. Balancing state and volunteer investment in biodiversity monitoring for the implementation of CBD indicators: A French example. Ecological Economics 69, 1580–1586 (2010).

57. Schmeller, D. S. et al. Building capacity in biodiversity monitoring at the global scale. Biodivers Conserv 26, 2765–2790 (2017).

58. IPBES. The Methodological Assessment Report on Scenarios and Models of Biodiversity and Ecosystem Services. https://zenodo.org/record/3235428 (2016) doi:10.5281/ZENODO.3235428.

59. Leung, B. & Gonzalez, A. Global monitoring for biodiversity: Uncertainty, risk, and power analyses to support trend change detection. Sci. Adv. 10, eadj1448 (2024).

60. Vaz, A. S. et al. The journey to monitoring ecosystem services: Are we there yet? Ecosystem Services 50, 101313 (2021).

61. CBD. Headline indicator D.3 on private funding: a background note. (2024).

62. Muñoz-García, M., Lago, A. & Scholz, H. A. Access and Benefit Sharing Indicators for the Kunming-Montreal Global Biodiversity Framework. 97 https://www.cbd.int/doc/c/6920/4e1e/8a6ba925279ea19033eb8ed2/sbstta-26-inf-12-en.pdf (2024).

63. Secretariat of the Convention on Biological Diversity. Nagoya Protocol on Access to Genetic Resources and the Fair and Equitable Sharing of Benefits Arising from their Utilization to the Convention on Biological Diversity. (2011).

64. Pereira, H. M. et al. Essential Biodiversity Variables. Science 339, 277–278 (2013).

65. Brummitt, N. et al. Taking stock of nature: Essential biodiversity variables explained. Biological Conservation 213, 252–255 (2017).

66. Balvanera, P. et al. Essential ecosystem service variables for monitoring progress towards sustainability. Current Opinion in Environmental Sustainability 54, 101152 (2022).

67. Mantyka-Pringle, C. S. et al. Bridging science and traditional knowledge to assess cumulative impacts of stressors on ecosystem health. Environment International 102, 125–137 (2017).

68. Brondízio, E. S. et al. Locally Based, Regionally Manifested, and Globally Relevant: Indigenous and Local Knowledge, Values, and Practices for Nature. Annu. Rev. Environ. Resour. 46, 481–509 (2021).

69. McElwee, P. et al. Working with Indigenous and local knowledge (ILK) in large scale ecological assessments: Reviewing the experience of the IPBES Global Assessment. Journal of Applied Ecology 57, 1666–1676 (2020).

70. Keith, D. A. et al. A function-based typology for Earth’s ecosystems. Nature 610, 513–518 (2022).

71. Economic and Social Council. Statistical Commission Report on the Fifty-Fifth Session. https://unstats.un.org/UNSDWebsite/statcom/session_55/documents/2024-36-FinalReport-E.pdf (2024).

72. SNBI & UNEP-WCMC. Mapping Biodiversity Priorities: A Practical Approach to Spatial Biodiversity Assessment and Prioritisation to Inform National Policy, Planning, Decisions and Action. (South African National Biodiveristy Institute, Pretoria, 2024).

73. GEO. Mapping the World’s Ecosystems for Action. (2024).

74. Berendsohn, W. et al. Biodiversity information platforms: From standards to interoperability. ZK 150, 71–87 (2011).

75. Navarro, L. M. et al. Monitoring biodiversity change through effective global coordination. Current Opinion in Environmental Sustainability 29, 158–169 (2017).

76. Gawich, M., Badr, A., Hegazy, A. & Ismail, H. A Methodology for Ontology Building. IJCA 56, 39–45 (2012).

77. Nieland, S., Kleinschmit, B. & Förster, M. Using ontological inference and hierarchical matchmaking to overcome semantic heterogeneity in remote sensing-based biodiversity monitoring. International Journal of Applied Earth Observation and Geoinformation 37, 133–141 (2015).

78. Griffith, J., et al. BON in a Box: An Open and Collaborative Platform for Biodiversity Monitoring, Indicator Calculation, and Reporting. Preprint at 10.32942/X2M320 (2024).

79. Walls, R. L. et al. Semantics in Support of Biodiversity Knowledge Discovery: An Introduction to the Biological Collections Ontology and Related Ontologies. PLoS ONE 9, e89606 (2014).

80. Kissling, W. D. et al. Towards global interoperability for supporting biodiversity research on essential biodiversity variables (EBVs). Biodiversity 16, 99–107 (2015).

81. Hardisty, A. R. et al. The Bari Manifesto: An interoperability framework for essential biodiversity variables. Ecological Informatics 49, 22–31 (2019).

82. Johnson, T. F. et al. Revealing uncertainty in the status of biodiversity change. Nature 628, 788–794 (2024).

83. Rowland, J. A., Bland, L. M., James, S. & Nicholson, E. A guide to representing variability and uncertainty in biodiversity indicators. Conservation Biology 35, 1669–1682 (2021).

84. Kühl, H. S. et al. Effective Biodiversity Monitoring Needs a Culture of Integration. One Earth 3, 462–474 (2020).

85. IPBES. Scoping report for a methodological assessment on monitoring biodiversity and nature’s contributions to people. (2023).

86. Scholes, R. J., Gill, M. J., Costello, M. J., Sarankatos, G. & Walters, M. Working in Networks to Make Biodiversity Data More Available. in The GEO Handbook on Biodiversity Observation Networks 1–18 (Springer International Publishing, 2017).

87. Gonzalez, A. et al. A global biodiversity observing system to unite monitoring and guide action. Nat Ecol Evol 7, 1947–1952 (2023).

88. WMO Integrated Global Observing System (WIGOS) | World Meteorological Organization. https://community.wmo.int/en/activity-areas/WIGOS (2024).

89. Pettorelli, N. et al. Time to integrate global climate change and biodiversity science policy agendas. Journal of Applied Ecology 58, 2384–2393 (2021).

90. García, F. C., Bestion, E., Warfield, R. & Yvon-Durocher, G. Changes in temperature alter the relationship between biodiversity and ecosystem functioning. Proc. Natl. Acad. Sci. U.S.A. 115, 10989–10994 (2018).

91. Pires, A. P. F. et al. Interactive effects of climate change and biodiversity loss on ecosystem functioning. Ecology 99, 1203–1213 (2018).

92. Levin, S. A. The Problem of Pattern and Scale in Ecology: The Robert H. MacArthur Award Lecture. Ecology 73, 1943–1967 (1992).

93. Mace, G. M., Norris, K. & Fitter, A. H. Biodiversity and ecosystem services: a multilayered relationship. Trends in Ecology & Evolution 27, 19–26 (2012).

94. Rounsevell, M. D. A., Dawson, T. P. & Harrison, P. A. A conceptual framework to assess the effects of environmental change on ecosystem services. Biodivers Conserv 19, 2823–2842 (2010).

95. Seddon, N. et al. Understanding the value and limits of nature-based solutions to climate change and other global challenges. Phil. Trans. R. Soc. B 375, 20190120 (2020).

96. Mori, A. S. et al. Biodiversity–productivity relationships are key to nature-based climate solutions. Nat. Clim. Chang. 11, 543–550 (2021).

97. Wassénius, E., Crona, B. & Quahe, S. Essential environmental impact variables: A means for transparent corporate sustainability reporting aligned with planetary boundaries. One Earth 7, 211–225 (2024).

98. CBD. Scientific and technical review of the traditional knowledge indicators and their suggested links with the headline, component and complementary indicators of the monitoring framework for the Kunming-Montreal Global Biodiversity Framework. (2024).

99. OHCHR. Human Rights Indicators: A Guide to Measurement and Implementation. (United Nations, 2013). doi:10.18356/58576336-en.

100. OHCHR. Guidance Note on a Human Rights-Based Approach to Data. (2018).

101. Ferrari, M. F., De Jong, C. & Belohrad, V. S. Community-based monitoring and information systems (CBMIS) in the context of the Convention on Biological Diversity (CBD). Biodiversity 16, 57–67 (2015).

102. Rovillos, D. R. & Dacquigan, A. E. L. The Indigenous Navigator: Human Rights, Sustainable Development Goals and the Indigenous Peoples in the Philippines. 64 https://indigenousnavigator.org/files/media/document/IN_Philippines_Report%20%28002%29.pdf (2024).

103. Reyes-García, V. et al. Recognizing Indigenous peoples’ and local communities’ rights and agency in the post-2020 Biodiversity Agenda. Ambio 51, 84–92 (2022).

104. Reyes-García, V. et al. The contributions of Indigenous Peoples and local communities to ecological restoration. Restoration Ecology 27, 3–8 (2019).

105. Lyver, P. O. B. et al. An indigenous community-based monitoring system for assessing forest health in New Zealand. Biodivers Conserv 26, 3183–3212 (2017).

106. Sullivan, J. J. & Molles, L. E. Biodiversity monitoring by community-based restoration groups in New Zealand. Ecological Management & Restoration 17, 210–217 (2016).

107. Ortega-Álvarez, R., Sánchez-González, L. A., Valera-Bermejo, A. & Berlanga-García, H. Community-Based Monitoring and Protected Areas: Towards an Inclusive Model. Sustainable Development 25, 200–212 (2017).

108. CBD. Review of implementation: progress in national target setting and updating of national biodiversity strategies and action plans. (2024).

109. Fisher, B. & Christopher, T. Poverty and biodiversity: Measuring the overlap of human poverty and the biodiversity hotspots. Ecological Economics 62, 93–101 (2007).

110. Flammer, C., Giroux, T. & Heal, G. Biodiversity Finance. w31022 http://www.nber.org/papers/w31022.pdf (2023) doi:10.3386/w31022.

111. WEF & PwC. Nature Risk Rising: Why the Crisis Engulfing Nature Matters for Business and the Economy. https://www3.weforum.org/docs/WEF_New_Nature_Economy_Report_2020.pdf (2020).

112. Smith, T. et al. Biodiversity means business: Reframing global biodiversity goals for the private sector. CONSERVATION LETTERS 13, e12690 (2020).

113. CBD. Decision adopted by the Conference of the Parties to the Convention on Biological Diversity at its tenth meeting. (2010).

114. ten Kate, K. & Laird, S. A. The Commercial Use of Biodiversity: Access to Genetic Resources and Benefit-Sharing. (Earthscan Publ, London, 2000).

115. Prathapan, K. D. et al. When the cure kills—CBD limits biodiversity research. Science 360, 1405–1406 (2018).

116. CBD. Guidance on using the indicators of the monitoring framework of the Kunming-Montreal Global Biodiversity Framework. (2024).

117. Rasoavahiny, L., Andrianarisata, M., Razafimpahanana, A. & Ratsifandrihamanana, A. N. Conducting an ecological gap analysis for the new Madagascar protected area system. Parks 17, 12–21 (2008).

118. Sumanapala, D. & Wolf, I. D. Recreational Ecology: A Review of Research and Gap Analysis. Environments 6, 81 (2019).

119. Drew, C. A. & Perera, A. H. Expert Knowledge as a Basis for Landscape Ecological Predictive Models. in Predictive Species and Habitat Modeling in Landscape Ecology (eds. Drew, C. A., Wiersma, Y. F. & Huettmann, F.) 229–248 (Springer New York, New York, NY, 2011). doi:10.1007/978-1-4419-7390-0_12.

120. Rodrigues, A. S. L. et al. Global Gap Analysis: Priority Regions for Expanding the Global Protected-Area Network. BioScience 54, 1092 (2004).

121. McBride, M. F. & Burgman, M. A. What Is Expert Knowledge, How Is Such Knowledge Gathered, and How Do We Use It to Address Questions in Landscape Ecology? in Expert Knowledge and Its Application in Landscape Ecology (eds. Perera, A. H., Drew, C. A. & Johnson, C. J.) 11–38 (Springer New York, New York, NY, 2012). doi:10.1007/978-1-4614-1034-8_2.

122. Knol, A. B., Slottje, P., Van Der Sluijs, J. P. & Lebret, E. The use of expert elicitation in environmental health impact assessment: a seven step procedure. Environ Health 9, 19 (2010).

123. Johnson, C. J., Drew, C. A. & Perera, A. H. Elicitation and Use of Expert Knowledge in Landscape Ecological Applications: A Synthesis. in Expert Knowledge and Its Application in Landscape Ecology (eds. Perera, A. H., Drew, C. A. & Johnson, C. J.) 279–299 (Springer New York, New York, NY, 2012). doi:10.1007/978-1-4614-1034-8_14.

124. Wizenried, A. Delphi Studies: The Value of Expert Opinion Bridging the Gap - Data to Knowledge. iasl 329–334 (2021) doi:10.29173/iasl8210.

125. Cousins, M. et al. Is scientific evidence enough? Using expert opinion to fill gaps in data in antimicrobial resistance research. PLoS ONE 18, e0290464 (2023).

126. Burgman, M. et al. Redefining expertise and improving ecological judgment: Ecological judgment. Conservation Letters 4, 81–87 (2011).

127. Kuhnert, P. M. Four case studies in using expert opinion to inform priors. Environmetrics 22, 662–674 (2011).

128. Hemming, V., Armstrong, N., Burgman, M. A. & Hanea, A. M. Improving expert forecasts in reliability: Application and evidence for structured elicitation protocols. Quality & Reliability Eng 36, 623–641 (2020).

129. Burgman, M. A. et al. Expert Status and Performance. PLoS ONE 6, e22998 (2011).

130. CBD. Monitoring framework for the Kunming-Montreal Global Biodiversity Framework. (2024).

131. CBD. Monitoring framework for the Kunming-Montreal Global Biodiversity Framework. (2024).

132. CBD. Monitoring framework for the Kunming-Montreal Global Biodiversity Framework. (2024).

133. Biodiversity Indicators Partnership. Guidance for national biodiversity indicator development and use. (2011).

134. Secretariat of the Convention on Biological Diversity. Convention on Biological Diversity: text and annexes. (2011).

135. CBD. Clearing House Mechanism Online Reporting Tool. https://ort.cbd.int/ (2024).

136. CBD. Mechanisms for planning, monitoring, reporting and review. (2024).

