## Supplementary material for "Assessing coverage of the Monitoring Framework of the Kunming-Montreal Global Biodiversity Framework and opportunities to fill gaps"

**Supplementary Table 1.** List of headline and binary indicators in the monitoring framework.

| Goal/Target | Indicator | Indicator name |
|---|---|---|
| Goal A, Target 1 | A.1 | Red List of Ecosystems |
| Goal A, Target 1 | A.2 | Extent of natural ecosystems |
| Goal A, Target 4 | A.3 | Red List Index (Sustainable Development Goal indicators 15.5.1) |
| Goal A, Target 4 | A.4 | The proportion of populations within species with an effective population size greater than 500 |
| Goal B, Target 11 | B.1 | Services provided by ecosystems |
| Goal B | B.b | Number of countries with policies or actions for implementing and monitoring the sustainable use of biodiversity and the maintenance and enhancement of nature's contributions to people, including ecosystem functions and services |
| Goal C, Target 13 | C.1 | Monetary benefits received in accordance with applicable internationally agreed access and benefit-sharing instruments |
| Goal C, Target 13 | C.2 | Non-monetary benefits arising from applicable international access and benefit-sharing instruments |
| Goal D, Target 19 | D.1 | International public funding, including official development assistance for conservation and sustainable use of biodiversity and ecosystems |
| Goal D, Target 19 | D.2 | Domestic public funding on conservation and sustainable use of biodiversity and ecosystems |
| Goal D, Target 19 | D.3 | Private funding (domestic and international) on conservation and sustainable use of biodiversity and ecosystems |
| Target 1 | 1.1 | Percentage of land and sea area covered by biodiversity-inclusive spatial plans |
| Target 1 | 1.b | Number of countries using participatory, integrated and biodiversity-inclusive spatial planning and/or effective management processes addressing land- and sea-use change to bring the loss of areas of high biodiversity importance close to zero by 2030 |
| Target 2 | 2.1 | Area under restoration |
| Target 3 | 3.1 | Coverage of protected areas and other effective area-based conservation measures |
| Target 5 | 5.1 | Proportion of fish stocks within biologically sustainable levels (Sustainable Development Goal indicator 14.4.1) |

| | | |
|---|---|---|
| Target 5 | 5.b | Number of countries with legal instruments or other policy frameworks to regulate trade in wild species |
| Target 6 | 6.1 | Rate of invasive alien species establishment |
| Target 6 | 6.b | Number of countries adopting relevant regulations, processes and measures to reduce the impact of invasive alien species |
| Target 7 | 7.1 | Index of coastal eutrophication potential (Sustainable Development Goal indicator 14.1.1 (a)) |
| Target 7 | 7.2 | Aggregated total applied toxicity |
| Target 8 | 8.b | Number of countries with policies to minimize the impact of climate change and ocean acidification on biodiversity and to minimize negative and foster positive impacts of climate action on biodiversity |
| Target 9 | 9.1 | Benefits from the sustainable use of wild species |
| Target 9 | 9.2 | Percentage of the population in traditional occupations |
| Target 9 | 9.b | Number or countries with policies to manage the use of wild species sustainably, providing social, economic and environmental benefits for people, and to protect and encourage customary sustainable use by indigenous peoples and local communities |
| Target 10 | 10.1 | Proportion of agricultural area under productive and sustainable agriculture (Sustainable Development Goal indicator 2.4.1) |
| Target 10 | 10.2 | Progress towards sustainable forest management (Sustainable Development Goal indicator 15.2.1) |
| Target 12 | 12.1 | Average share of the built-up area of cities that is green or blue space for public use for all |
| Target 12 | 12.b | Number of countries with biodiversity-inclusive urban planning referring to green or blue urban spaces |
| Target 13 | 13.b | Number of countries that have taken effective legal, policy, administrative and capacity- building measures at all levels, as appropriate, to ensure the fair and equitable sharing of benefits from the utilization of genetic resources and from digital sequence information on genetic resources, as well as traditional knowledge associated with genetic resources. |
| Target 14 | 14.b | Number of countries integrating biodiversity and its multiple values into policies, regulations, planning, development processes, poverty eradication strategies and, as appropriate, national accounts, within and across all levels and across all sectors, and progressively aligning all relevant public and private activities and fiscal and financial flows with the goals and targets of the Framework |
| Target 15 | 15.1 | Number of companies disclosing their biodiversity-related risks, dependencies and impacts |
| Target 15 | 15.b | Number of countries with legal, administrative or policy measures aimed at encouraging and enabling business and financial institutions, and in particular for large and transnational companies |

| | | |
|---|---|---|
| | | and financial institutions, to progressively reduce their negative impacts on biodiversity, increase their positive impacts, reduce their biodiversity-related risks and promote actions to ensure sustainable patterns of production |
| Target 16 | 16.b | Number of countries developing, adopting or implementing policy instruments aimed at encouraging and enabling people to make sustainable consumption choices |
| Target 17 | 17.b | Number of countries that have taken action to implement biosafety measures as set out in Article 8(g) of the Convention and measures for the handling of biotechnology and the distribution of its benefits as set out in Article 19 |
| Target 18 | 18.1 | Positive incentives in place to promote biodiversity conservation and sustainable use |
| Target 18 | 18.2 | Value of subsidies and other incentives harmful to biodiversity |
| Target 20 | 20.b | Number of countries that have taken significant action to strengthen capacity-building and development and access to and transfer of technology, and to promote the development of and access to innovation and technical and scientific cooperation |
| Target 21 | 21.1 | Indicator on biodiversity information for monitoring the Kunming-Montreal Global Biodiversity Framework |
| Target 22 | 22.1 | Land-use change and land tenure in the traditional territories of indigenous peoples and local communities |
| Target 22 | 22.b | Number of countries taking action towards the full, equitable, inclusive, effective and gender-responsive representation and participation, in decision-making, and access to justice and information related to biodiversity by indigenous peoples and local communities, respecting their cultures and their rights over lands, territories, resources, and traditional knowledge, as well as by, women, and girls, children and youth, and persons with disabilities and the full protection of environmental human rights defenders |
| Target 23 | 23.b | Number of countries with legal, administrative or policy frameworks, inter alia, the Gender Plan of Action (2023–2030), to ensure that all women and girls have equal opportunity and capacity to contribute to the three objectives of the Convention, including by ensuring women's equal rights and access to land and natural resources |

**Supplementary Table 2.** List of distinct elements identified in the text of the GBF that require monitoring for a holistic progress tracking. Here, the text was taken from the Goals and Targets alone, without the corresponding and relevant considerations set out in Section C of the GBF, which are to be considered cross-cutting.

| Goal/Target | Element |
|---|---|

| | |
|---|---|
| Goal A | A (i) The **integrity** of all ecosystems is maintained, enhanced, or restored |
| | A (ii) The **connectivity** of all ecosystems is maintained, enhanced, or restored |
| | A (iii) The **resilience** of all ecosystems is maintained, enhanced, or restored |
| | A (iv) The **area of natural ecosystems** is substantially increased by 2050 |
| | A (v) Human induced **extinction of known threatened species** is halted |
| | A (vi) By 2050, the **extinction rate** of all species is reduced tenfold |
| | A (vii) By 2050, the **extinction risk** of all species is reduced tenfold |
| | A (viii) By 2050**,** the **abundance of native wild species** is increased to healthy and resilient levels |
| | A (ix) The **genetic diversity within populations of wild species** is maintained, safeguarding their adaptive potential |
| | A (x) The **genetic diversity within populations of** domestic **species** is maintained, safeguarding their adaptive potential |
| Goal B | B (i) Biodiversity is **sustainably used and managed** |
| | B (ii) **Nature's contributions to people**, including ecosystem functions and services, are **valued** |
| | B (iii) **Nature's contributions to people**, including ecosystem functions and services, are **maintained** |
| | B (iv) **Nature's contributions to people**, including ecosystem functions and services, are **enhanced** |
| | B (v) **Nature's contributions to people** currently in decline are **restored** |
| | B (vi) **supporting the achievement of sustainable development for the benefit of present and future generations by 2050.** |
| Goal C | C (i) The **monetary benefits** from the **utilization of genetic resources are shared fairly and equitably,** including, as appropriate with indigenous peoples and local communities, and substantially increased by 2050 |
| | C (ii) The **monetary benefits** from the **utilization of digital sequence information are shared fairly and equitably**, including, as appropriate with indigenous peoples and local communities, and substantially increased by 2050 |

| | |
|---|---|
| | C (iii) The **monetary benefits** from the **utilization of traditional knowledge associated with genetic resources are shared fairly and equitably,** including, as appropriate with indigenous peoples and local communities, and substantially increased by 2050 |
| | C (iv) The **non-monetary benefits** from the **utilization of genetic resources are shared fairly and equitably**, including, as appropriate with indigenous peoples and local communities, and substantially increased by 2050 |
| | C (v) The **non-monetary benefits** from the **utilization of digital sequence information are shared fairly and equitably,** including, as appropriate with indigenous peoples and local communities, and substantially increased by 2050 |
| | C (vi) The **non-monetary benefits** from the **utilization of traditional knowledge associated with genetic resources are shared fairly and equitably**, including, as appropriate with indigenous peoples and local communities, and substantially increased by 2050 |
| | C (vii) **Traditional knowledge** associated with genetic resources is appropriately pro**tected** |
| Goal D | D (i) **Adequate financial resources secured** |
| | D (ii) **Adequate capacity-building secured** |
| | D (iii) **Adequate technical and scientific cooperation secured** |
| | D (iv) **Access to and transfer of technology secured** |
| | D (v) **Means of implementation are equitably accessible to all Parties** |
| Target 1 | 1a. **All areas are under spatial planning and/or effective management processes addressing land/sea use change** |
| | 1b. **Spatial planning is participatory** |
| | 1c. **Spatial planning is integrated** |
| | 1d. **Spatial planning is biodiversity-inclusive** |
| | 1e. **Spatial planning/management processes bring the loss of areas of high biodiversity importance, including ecosystems of high ecological integrity, close to zero by 2030** |
| | 1f. **Spatial planning/management processes respect the rights of IPLCs** |

| | |
|---|---|
| Target 2 | 2a. Ensure that by 2030, **at least 30 per cent of areas of degraded terrestrial ecosystems are under restoration** |
| | 2b. Ensure that by 2030, **at least 30 per cent of degraded inland water ecosystems are under restoration** |
| | 2c. Ensure that by 2030, **at least 30 per cent of degraded marine and coastal ecosystems are under restoration** |
| | 2d. Restoration is **effective**, **enhancing biodiversity** |
| | 2e. Restoration is **effective, enhancing ecosystem functions and services** |
| | 2f. Restoration is **effective**, **enhancing ecological integrity and connectivity** |
| Target 3 | 3a. Ensure and enable that by 2030 **at least 30 per cent of terrestrial and inland water areas are included in systems of protected areas and OECMs, recognizing indigenous and traditional territories where applicable** |
| | 3b. Ensure and enable that by 2030 **at least 30 per cent of marine and coastal areas are included in systems of protected areas and OECMs, recognizing indigenous and traditional territories where applicable** |
| | 3c. Ensure and enable that by 2030, **areas of particular importance for biodiversity are included in systems of protected areas and OECMs, recognizing indigenous and traditional territories where applicable** |
| | 3d. Ensure and enable that by 2030, **areas of particular importance for ecosystem functions and services are included in systems of protected areas and OECMs, recognizing indigenous and traditional territories where applicable** |
| | 3e. Systems of protected areas and OECMS, recognizing indigenous and traditional territories where applicable, **effectively manage and conserve** terrestrial, inland water, marine and coastal areas |
| | 3f. Systems of protected areas and OECMS, recognizing indigenous and traditional territories where applicable, are **ecologically representative** |
| | 3g. Systems of protected areas and OECMS, recognizing indigenous and traditional territories where applicable, are **well connected** |
| | 3h. Systems of protected areas and OECMS, recognizing indigenous and traditional territories where applicable, are **equitably governed** |
| | 3i. Systems of protected areas and OECMS, recognizing indigenous and traditional territories where applicable, are **integrated into wider landscapes, seascapes and the ocean** |

| | |
|---|---|
| | 3j. Ensure that any **sustainable use,** where appropriate in such areas i**s fully consistent with conservation outcomes** |
| | 3k. Systems of protected areas and OECMs **recognize and respect the rights of IPLCs, including over their traditional territories** |
| Target 4 | 4a. **Ensure urgent management actions including through *in situ* and *ex situ* conservation and sustainable management practices** |
| | 4b. **Halt human-induced extinction of known threatened species** |
| | 4c. Enable the **recovery and conservation of species, in particular threatened species** |
| | 4d. Maintain and restore the **genetic diversity within and between populations of native, wild species** to maintain their adaptive potential |
| | 4e. Maintain and restore the **genetic diversity within and between populations of domestic species** to maintain their adaptive potential |
| | 4f. Effectively manage **human-wildlife interactions to minimize human-wildlife conflicts** for coexistence |
| Target 5 | 5a. Ensure that the **use and harvesting of wild species is sustainable** preventing overexploitation, minimizing impacts on non-target species and ecosystems |
| | 5b. Ensure that the **trade in wild species** is sustainable preventing overexploitation, minimizing impacts on non-target species and ecosystems |
| | 5c. Ensure that the use, harvesting and trade of wild species is **safe, reducing the risk of pathogen spillover** |
| | 5d. Ensure that the use, harvesting and trade of wild species is **legal** |
| | 5e. **Apply the ecosystem approach** in addressing sustainable, safe and legal use, harvesting and trade of wild species |
| | 5f. **Respect and protect customary sustainable use by IPLCs** in addressing sustainable, safe and legal use, harvesting and trade of wild species |
| Target 6 | 6a. Eliminate, minimize, reduce and or mitigate **the impacts of invasive alien species on biodiversity** |
| | 6b. Eliminate, minimize, reduce and or mitigate the **impacts of invasive alien species on ecosystem services** |
| | 6c. **Identify pathways** of the introduction of invasive alien species |
| | 6d. **Manage pathways** of the introduction of invasive alien species |

| | |
|---|---|
| | 6e. **Prevent the introduction and establishment of priority invasive alien species** |
| | 6f. **Reduce the rates of introduction and establishment of other known or potential invasive alien species by at least 50 per cent** by 2030 |
| | 6g. **Eradicate or control** invasive alien species, **especially on priority sites, such as islands** |
| Target 7 | 7a. **Reduce pollution risks and the negative impact of pollution** from all sources, by 2030, **to levels that are not harmful to biodiversity** considering cumulative effects |
| | 7b. **Reduce pollution risks and the negative impact of pollution** from all sources, by 2030, **to levels that are not harmful to ecosystem functions and services**, considering cumulative effects |
| | 7c. **Reduce excess nutrients lost to the environment by at least half** including through more efficient nutrient cycling and use |
| | 7d. **Reduce the overall risk from pesticides by at least half** including through integrated pest management, based on science, taking into account food security and livelihoods |
| | 7e. **Reduce the overall risk from highly hazardous chemicals by at least half** |
| | 7f. Work towards **eliminating plastic pollution.** |
| Target 8 | 8a. **Minimize the impact of climate change on biodiversity**, **and increase its resilience,** through mitigation, adaptation and disaster risk reduction actions |
| | 8b. **Minimize the impact of ocean acidification on biodiversity, and increase its resilience,** through mitigation, adaptation and disaster risk reduction actions |
| | 8c. Apply **nature-based solutions** (in minimizing climate change/ocean acidification impacts and increasing resilience) |
| | 8d. Apply the **ecosystem-based approach** (in minimizing climate change/ocean acidification impacts and increasing resilience) |
| | 8e. Minimize **negative impacts of climate action on biodiversity** |
| | 8f. Foster **positive impacts of climate action on biodiversity** |
| Target 9 | 9a. Ensure that the **management and use of wild species are sustainable**, thereby providing **social benefits for people** |

| | |
|---|---|
| | 9b. Ensure that the **management and use of wild species are sustainable**, thereby providing **economic benefits for people** |
| | 9c. Ensure that the **management and use of wild species are sustainable**, thereby providing **environmental benefits for people** |
| | 9d. Sustainable management and use of wild species provides benefits for **people in vulnerable situations and those most dependent on biodiversity** |
| | 9e. Ensure benefits to people through **sustainable biodiversity-based activities, products and services that enhance biodiversity** |
| | 9f. Protect and encourage **customary sustainable use by IPLCs** |
| Target 10 | 10a. Ensure that **areas under agriculture are managed sustainably** |
| | 10b. Ensure that **areas under aquaculture are managed sustainably** |
| | 10c. Ensure that **areas under fisheries are managed sustainably** |
| | 10d. Ensure that **areas under forestry are managed sustainably** |
| | 10e. Ensure sustainable management of productive sectors through a **substantial increase of the application of biodiversity friendly practices, such as sustainable intensification, agroecological and other innovative approaches** |
| | 10f. Ensure that sustainable management of productive sectors contributes to the **resilience and long-term efficiency and productivity of these production systems** |
| | 10g. Ensure that sustainable management of productive sectors contributes to **food security** |
| | 10h. Ensure that sustainable management of productive sectors contributes to **conserving and restoring biodiversity** |
| | 10i. Ensure that sustainable management of productive sectors contributes to **maintaining nature's contributions to people, including ecosystem functions and services** |
| Target 11 | 11a. **Restore, maintain and enhance nature's contributions to people**, including ecosystem functions and services **through nature-based solutions** for the benefit of all people and nature |
| | 11b. **Restore, maintain and enhance nature's contributions to people**, including ecosystem functions and services **through ecosystem-based approaches** for the benefit of all people and nature |
| | 11c. Restore, maintain and enhance **regulation of air, water and climate** |
| | 11d. Restore, maintain and enhance **soil health** |

| | |
|---|---|
| | 11e. Restore, maintain and enhance **pollination** |
| | 11f. Restore, maintain and enhance **reduction of disease risk** |
| | 11g. Restore, maintain and enhance **protection from natural hazards and disasters** |
| Target 12 | 12a. Significantly increase the **area of green spaces in urban and densely populated areas** |
| | 12b. Significantly increase the **area of blue spaces in urban and densely populated areas** |
| | 12c. Significantly increase the **quality of green and blue spaces in urban and densely populated areas** |
| | 12d. Significantly increase the **connectivity of green and blue spaces in urban and densely populated areas** |
| | 12e. Significantly increase the **access to and benefits from green and blue spaces in urban and densely populated areas** |
| | 12f. **Mainstream the conservation and sustainable use of biodiversity in urban and densely populated areas** |
| | 12g. Ensure **biodiversity-inclusive urban planning** contributing to inclusive and sustainable urbanization and to the provision of ecosystem functions and services. |
| | 12h. **Enhance native biodiversity in urban and densely populated areas** |
| | 12i. **Improve human health and well-being and connectedness to nature,** through biodiversity-inclusive urban planning |
| Target 13 | 13a. Take effective **legal, policy and administrative measures** at all levels, as appropriate, to **ensure the fair and equitable sharing of benefits that arise from the utilization of genetic resources** |
| | 13b. Take effective **capacity building measures** at all levels, as appropriate, **to ensure the fair and equitable sharing of benefits that arise from the utilization of genetic resources** |
| | 13c. Take effective **legal, policy and administrative** measures at all levels, as appropriate, to **ensure the fair and equitable sharing of benefits that arise from the utilization of digital sequence information on genetic resources** |
| | 13d. Take effective **capacity building measures** at all levels, as appropriate, to **ensure the fair and equitable sharing of benefits that arise from the utilization of digital sequence information on genetic resources** |
| | 13e. Take effective **legal, policy and administrative measures** at all levels, as appropriate, to **ensure the fair and equitable sharing of benefits that arise from traditional knowledge associated with genetic resources** |

| | |
|---|---|
| | 13f. Take effective **capacity building measures** at all levels, as appropriate, to **ensure the fair and equitable sharing of benefits that arise from traditional knowledge associated with genetic resources** |
| | 13g. **Facilitate appropriate access to genetic resources** |
| | 13h. By 2030, facilitate a **significant increase of the benefits shared**, in accordance with applicable international access and benefit-sharing instruments |
| Target 14 | 14a. **Ensure the full integration of biodiversity and its multiple values** into **policies, regulations, planning and development processes** |
| | 14b. **Ensure the full integration of biodiversity and its multiple values** into **poverty eradication strategies** |
| | 14c. **Ensure the full integration of biodiversity and its multiple values** into **strategic environmental assessments and environmental impact assessments** |
| | 14d. **Ensure the full integration of biodiversity and its multiple values** into **national accounting,** as appropriate |
| | 14e. **Ensure the full integration of biodiversity and its multiple values within and across all levels of government** |
| | 14f. **Ensure the full integration of biodiversity and its multiple values across all sectors, in particular those with significant impacts on biodiversity** |
| | 14g. Progressively **align all relevant public and private activities, and fiscal and financial flows** with the goals and targets of this framework |
| Target 15 | 15a. Take legal, administrative or policy measures to **encourage and enable business,** and in particular to ensure that large and transnational companies and financial institutions, along their operations, supply and value chains and portfolios, regularly **monitor, assess and transparently disclose their risks relating to biodiversity** |
| | 15b. Take legal, administrative or policy measures to **encourage and enable business**, and in particular to ensure that large and transnational companies and financial institutions, along their operations, supply and value chains and portfolios, regularly **monitor, assess and transparently disclose their dependencies on biodiversity** |
| | 15c. Take legal, administrative or policy measures to **encourage and enable business**, and in particular to ensure that large and transnational companies and financial institutions, along their operations, supply and value chains and portfolios, regularly **monitor, assess and transparently disclose their impacts on biodiversity** |

| | |
|---|---|
| | 15d. Take legal, administrative or policy measures to **encourage and enable business**, and in particular to ensure that large and transnational companies and financial institutions, **provide information needed to consumers to promote sustainable consumption patterns** |
| | 15e.Take legal, administrative or policy measures to **encourage and enable business**, and in particular to ensure that large and transnational companies and financial institutions, **report on compliance with access and benefit-sharing regulations and measures, as applicable** |
| Target 16 | 16a. Ensure that **people are encouraged and enabled to make sustainable consumption choices by establishing supportive policy, legislative or regulatory frameworks** |
| | 16b. Ensure that **people are encouraged and enabled to make sustainable consumption choices by improving education and access to relevant and accurate information and alternatives** |
| | 16c. By 2030, **reduce the global footprint of consumption in an equitable manner** in order for all people to live well in harmony with Mother Earth |
| | 16d. By 2030, **halve global food waste** |
| | 16e. By 2030, **significantly reduce overconsumption** |
| | 16f. By 2030, **substantially reduce waste generation** |
| Target 17 | 17a. **Establish**, in all countries, **biosafety measures as set out in Article 8(g)** of the Convention on Biological Diversity |
| | 17b. **Strengthen capacity for**, in all countries, **biosafety measures as set out in Article 8(g)** of the Convention on Biological Diversity |
| | 17c. **Implement**, in all countries, **biosafety measures as set out in Article 8(g)** of the Convention on Biological Diversity |
| | 17d. **Establish**, in all countries, **measures for the handling of biotechnology and distribution of its benefits as set out in Article 19** of the Convention. |
| | 17e. **Strengthen capacity for**, in all countries, **measures for the handling of biotechnology and distribution of its benefits as set out in Article 19** of the Convention. |
| | 17f. **Implement, in all countries, measures for the handling of biotechnology and distribution of its benefits as set out in Article 19** of the Convention. |
| Target 18 | 18a. **Identify by 2025 incentives, including subsidies, harmful for biodiversity** |

| | |
|---|---|
| | 18b. **Eliminate incentives, including subsidies, harmful for biodiversity**, in a proportionate, just, fair, effective and equitable way |
| | 18c. **Phase out incentives, including subsidies, harmful for biodiversity**, in a proportionate, just, fair, effective and equitable way |
| | 18d. **Reform incentives, including subsidies, harmful for biodiversity**, in a proportionate, just, fair, effective and equitable way |
| | 18e. **Substantially and progressively reduce incentives, including subsidies, harmful for biodiversity, by at least $500 billion per year by 2030, starting with the most harmful incentives** |
| | 18f. **Scale up positive incentives for the conservation and sustainable use of biodiversity**. |
| Target 19 | 19a. **Increasing total biodiversity related international financial resources from developed countries and from countries** that voluntarily assume obligations of developed country Parties, to developing countries **to at least US$ 20 billion per year by 2025, and to at least US$ 30 billion per year by 2030.** |
| | 19b. **Significantly increasing domestic resource mobilization**, facilitated by the preparation and implementation of national biodiversity finance plans or similar instruments |
| | 19c. **Leveraging private finance, promoting blended finance, implementing strategies for raising new and additional resources, and encouraging the private sector to invest in biodiversity** |
| | 19d. **Stimulating innovative schemes such as payment for ecosystem services, green bonds, biodiversity offsets and credits, benefit-sharing mechanisms, with environmental and social safeguards.** |
| | 19e. **Optimizing co-benefits and synergies of finance** targeting the biodiversity and climate crises |
| | 19f. **Enhancing the role of collective actions, including by IPLCs, Mother Earth centric actions and non-market-based approaches** including community based natural resource management and civil society cooperation and solidarity aimed at the conservation of biodiversity |
| | 19g. **Enhancing the effectiveness, efficiency and transparency of resource provision and use** |
| Target 20 | 20a. **Strengthen capacity-building and development to meet the needs for effective implementation,** particularly in developing countries |
| | 20b. **Strengthen access to and transfer of technology to meet the needs for effective implementation,** particularly in developing countries |

| | |
|---|---|
| | 20c. **Promote development of and access to innovation to meet the needs for effective implementation**, particularly in developing countries |
| | 20d. **Promote technical and scientific cooperation, including through South-South, North-South and triangular cooperation, to meet the needs for effective implementation**, particularly in developing countries |
| | 20e. **Foster joint technology development for the conservation and sustainable use of biodiversity** |
| | 20f. **Foster joint scientific research programmes for the conservation and sustainable use of biodiversity** |
| | 20g. Strengthen scientific research and **monitoring capacities**, commensurate with the ambition of the goals and targets of the Framework. |
| Target 21 | 21a. **Ensure that the best available data is accessible to decision-makers, practitioners and the public** to guide effective and equitable governance, integrated and participatory management of biodiversity |
| | 21b. **Ensure that the best available information and knowledge is accessible to decision-makers, practitioners and the public** to guide effective and equitable governance, integrated and participatory management of biodiversity |
| | 21c. Strengthen **communication, awareness-raising and education** |
| | 21d. Strengthen **monitoring** |
| | 21e. Strengthen **research** |
| | 21f. Strengthen **knowledge management** |
| | 21g. (Ensure that) **traditional knowledge, innovations, practices and technologies of indigenous peoples and local communities** (are) **only be accessed with their free, prior and informed consent, in accordance with national legislation.** |
| Target 22 | 22a. **Ensure the full, equitable, inclusive, effective and gender-responsive representation and participation in decision-making related to biodiversity by IPLCs**, respecting their cultures and their rights over lands, territories, resources, and traditional knowledge |
| | 22b. **Ensure access to justice related to biodiversity by IPLCs**, respecting their cultures and their rights over lands, territories, resources, and traditional knowledge |
| | 22c.**Ensure access to information related to biodiversity by IPLCs**, respecting their cultures and their rights over lands, territories, resources, and traditional knowledge |
| | 22d. **Ensure the full, equitable, inclusive, effective and gender-responsive representation and participation in decision-making related to biodiversity by women and girls** |

| | |
|---|---|
| | 22e. **Ensure access to justice related to biodiversity by women and girls** |
| | 22f. **Ensure access to information related to biodiversity by women and girls** |
| | 22g. **Ensure the full, equitable, inclusive, effective and gender-responsive representation and participation in decision-making related to biodiversity by children and youth** |
| | 22h. **Ensure access to justice related to biodiversity by children and youth** |
| | 22i. **Ensure access to information related to biodiversity by children and youth** |
| | 22j. **Ensure the full, equitable, inclusive, effective and gender-responsive representation and participation in decision-making related to biodiversity by persons with disabilities** |
| | 22k **Ensure access to justice related to biodiversity by persons with disabilities** |
| | 22l. **Ensure access to information related to biodiversity by persons with disabilities** |
| | 22m. **Ensure the full protection of environmental human rights defenders.** |
| | 22n. **respecting their cultures and their rights over lands, territories, resources, and traditional knowledge** |
| Target 23 | 23a. **Ensure gender equality in the implementation of the Framework through a gender-responsive approach**, where all women and girls have equal opportunity and capacity to contribute to the three objectives of the Convention |
| | 23b. Recognize the **equal rights of women and girls to land and natural resources** |
| | 23c. Recognize the **equal access of women and girls to land and natural resources** |
| | 23d. Recognize **full, equitable, meaningful, and informed participation and leadership of women and girls at all levels of action, engagement, policy and decision-making related to biodiversity.** |

**Supplementary Table 3.** Number of distinct elements identified in the text of each goal and target.

| | Elements |
|---|---|
| Goal A | 10 |
| Goal B | 6 |
| Goal C | 7 |
| Goal D | 5 |
| Target 1 | 6 |
| Target 2 | 6 |
| Target 3 | 11 |

| | Elements |
|---|---|
| Target 4 | 6 |
| Target 5 | 6 |
| Target 6 | 7 |
| Target 7 | 6 |
| Target 8 | 6 |
| Target 9 | 6 |
| Target 10 | 9 |
| Target 11 | 7 |
| Target 12 | 9 |
| Target 13 | 8 |
| Target 14 | 7 |
| Target 15 | 5 |
| Target 16 | 6 |
| Target 17 | 6 |
| Target 18 | 6 |
| Target 19 | 7 |
| Target 20 | 7 |
| Target 21 | 7 |
| Target 22 | 14 |
| Target 23 | 4 |

**Supplementary Table 4.** Overall scores of coverage for each element by its headline and binary indicator(s). There are complete gaps in coverage where no indicators are listed and no score is given.

| Element | Indicator(s) | Score |
|---|---|---|
| A(i) | A.1 | Partially |
| A(ii) | A.1 | Partially |
| A(iii) | A.1 | Covered |
| A(iv) | A.1, A.2 | Covered |
| A(v) | A.3 | Partially |
| A(vi) | A.3 | Partially |
| A(vii) | A.4 | Covered |
| A(viii) | | |
| A(ix) | A.4 | Covered |
| A(x) | A.4 | Partially |
| B(i) | B.1, B.b | Partially |
| B(ii) | | |
| B(iii) | B.1, B.b | Covered |
| B(iv) | B.1, B.b | Covered |
| B(v) | B.b | Partially |

| | | |
|---|---|---|
| B(vi) | | |
| C(i) | C.1 | Potentially |
| C(ii) | C.1 | Potentially |
| C(iii) | C.1 | Potentially |
| C(iv) | C.2 | Potentially |
| C(v) | C.2 | Potentially |
| C(vi) | C.2 | Potentially |
| C(vii) | | |
| D(i) | D.1, D.2, D.3 | Covered |
| D(ii) | | |
| D(iii) | | |
| D(iv) | | |
| D(v) | | |
| 1a | 1.b | Covered |
| 1b | 1.b | Covered |
| 1c | 1.b | Partially |
| 1d | 1.b | Covered |
| 1e | A.1, A.2 | Partially |
| 1f | | |
| 2a | 2.2 | Partially |
| 2b | 2.2 | Partially |
| 2c | 2.2 | Partially |
| 2d | | |
| 2e | | |
| 2f | | |
| 3a | 3.1 | Covered |
| 3b | 3.1 | Covered |
| 3c | 3.1 | Covered |
| 3d | | |
| 3e | 3.1 | Covered |
| 3f | 3.1 | Covered |
| 3g | | |
| 3h | 3.1 | Partially |

| | | |
|---|---|---|
| 3i | 3.1 | Partially |
| 3j | | |
| 3k | | |
| 4a | | |
| 4b | A.3 | Partially |
| 4c | A.3 | Partially |
| 4d | A.4 | Partially |
| 4e | A.4 | Partially |
| 4f | | |
| 5a | 5.1 | Partially |
| 5b | | |
| 5c | | |
| 5d | | |
| 5e | | |
| 5f | | |
| 6a | 6.1, 6.b | Partially |
| 6b | 6.b | Potentially |
| 6c | 6.1 | Potentially |
| 6d | | |
| 6e | 6.1, 6.b | Partially |
| 6f | 6.1 | Covered |
| 6g | 6.b | Partially |
| 7a | 7.1, 7.2 | Partially |
| 7b | 7.1, 7.2 | Partially |
| 7c | 7.1 | Partially |
| 7d | 7.2 | Partially |
| 7e | 7.2 | Partially |
| 7f | | |
| 8a | 8.b | Partially |
| 8b | 8.b | Partially |
| 8c | 8.b | Covered |
| 8d | 8.b | Covered |
| 8e | 8.b | Partially |

| | | |
|---|---|---|
| 8f | 8.b | Partially |
| 9a | 9.b | Partially |
| 9b | 9.b | Partially |
| 9c | 9.b | Partially |
| 9d | 9.2 | Partially |
| 9e | 9.2, 9.b | Partially |
| 9f | 9.2, 9.b | Covered |
| 10a | 10.1 | Covered |
| 10b | | |
| 10c | 5.1 | Partially |
| 10d | 10.2 | Covered |
| 10e | 10.1, 10.2 | Partially |
| 10f | 10.1, 10.2 | Partially |
| 10g | 10.1 | Covered |
| 10h | 10.1, 10.2 | Partially |
| 10i | 10.1, 10.2 | Partially |
| 11a | | |
| 11b | | |
| 11c | B.1 | Potentially |
| 11d | B.1 | Potentially |
| 11e | B.1 | Potentially |
| 11f | B.1 | Potentially |
| 11g | B.1 | Potentially |
| 12a | 12.1 | Potentially |
| 12b | | |
| 12c | | |
| 12d | | |
| 12e | 12.1 | Partially |
| 12f | | |
| 12g | 12.b | Covered |
| 12h | | |
| 12i | | |
| 13a | 13.b | Covered |

| | | |
|---|---|---|
| 13b | C.2 | Potentially |
| 13c | C.1, C.2, 13.b | Covered |
| 13d | | |
| 13e | C.1, C.2, 13.b | Covered |
| 13f | C.2 | Potentially |
| 13g | | |
| 13h | C.1, C.2 | Potentially |
| 14a | 14.b | Partially |
| 14b | 14.b | Partially |
| 14c | 14.b | Partially |
| 14d | 14.b | Partially |
| 14e | 14.b | Partially |
| 14f | 14.b | Partially |
| 14g | 14.b | Partially |
| 15a | 15.1, 15.b | Covered |
| 15b | 15.1, 15.b | Covered |
| 15c | 15.1, 15.b | Covered |
| 15d | 15.b | Covered |
| 15e | 15.b | Covered |
| 16a | 16.b | Partially |
| 16b | 16.b | Partially |
| 16c | 16.b | Partially |
| 16d | | |
| 16e | 16.b | Potentially |
| 16f | 16.b | Potentially |
| 17a | 17.b | Covered |
| 17b | 17.b | |
| 17c | 17.b | Covered |
| 17d | 17.b | Covered |
| 17e | 17.b | Partially |
| 17f | 17.b | Covered |
| 18a | 18.2 | Covered |
| 18b | | |

| | | |
|---|---|---|
| 18c | | |
| 18d | | |
| 18e | 18.2 | Covered |
| 18f | 18.1 | Covered |
| 19a | D.1 | Covered |
| 19b | D.2 | Covered |
| 19c | D.3 | Covered |
| 19d | D.3 | Covered |
| 19e | | |
| 19f | | |
| 19g | | |
| 20a | 20.b | Partially |
| 20b | 20.b | Partially |
| 20c | 20.b | Partially |
| 20d | 20.b | Partially |
| 20e | 20.b | Partially |
| 20f | 20.b | Partially |
| 20g | 20.b | Partially |
| 21a | 21.1 | Potentially |
| 21b | 21.1 | Potentially |
| 21c | | |
| 21d | 21.1 | Potentially |
| 21e | 21.1 | Potentially |
| 21f | 21.1 | Potentially |
| 21g | 21.1 | Potentially |
| 22a | 22.b | Partially |
| 22b | 22.b | Partially |
| 22c | 22.b | Partially |
| 22d | 22.b | Partially |
| 22e | 22.b | Partially |
| 22f | 22.b | Partially |
| 22g | 22.b | Partially |
| 22h | 22.b | Partially |

| | | |
|---|---|---|
| 22i | 22.b | Partially |
| 22j | 22.b | Partially |
| 22k | 22.b | Partially |
| 22l | 22.b | Partially |
| 22m | 22.b | Partially |
| 22n | 22.b | Partially |
| 23a | 23.b | Partially |
| 23b | 23.b | Partially |
| 23c | 23.b | Partially |
| 23d | 23.b | Partially |

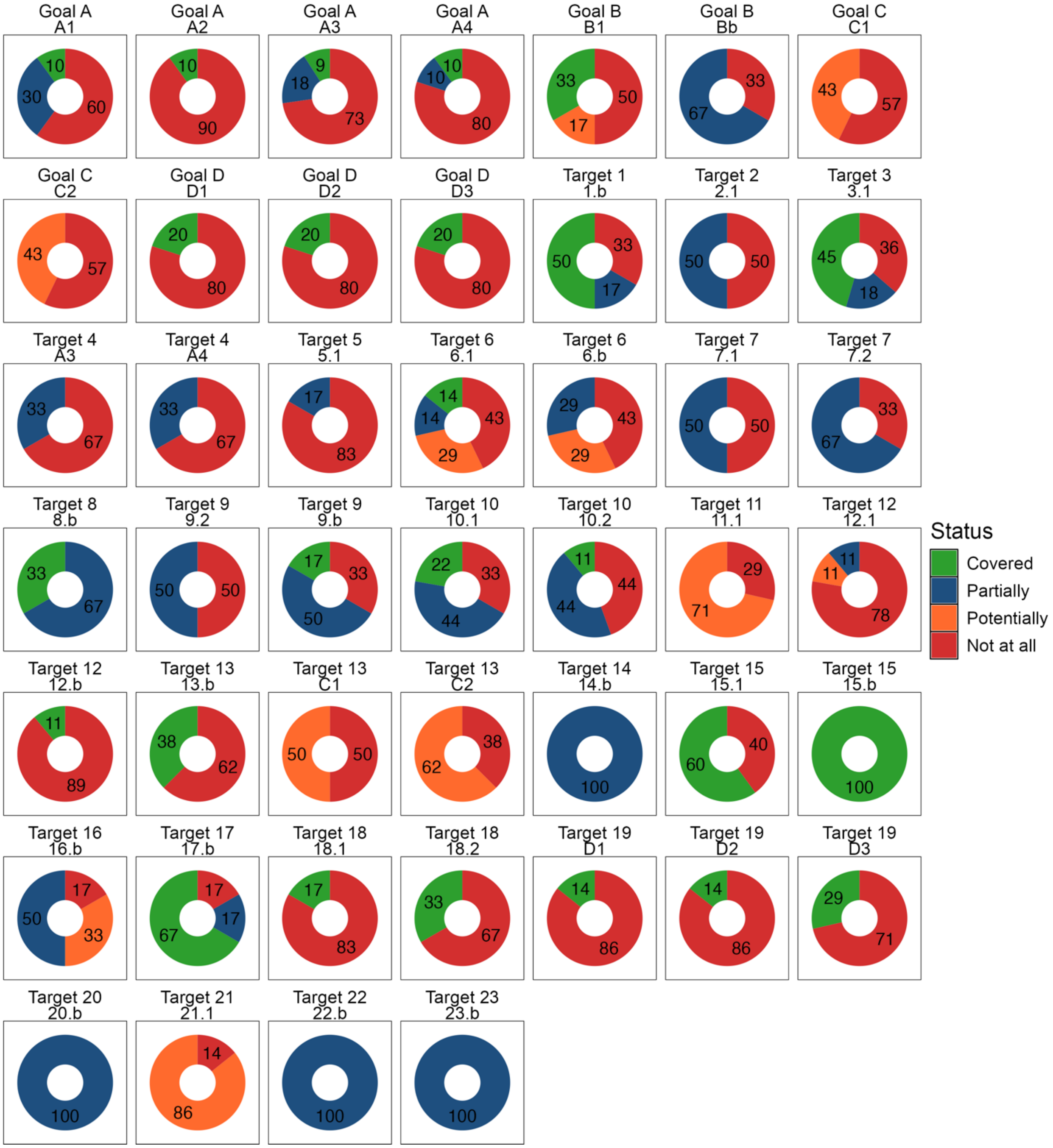


**Supplementary Figure 1.** Coverage of elements of each of the Goals and Targets of the GBF by their respective headline and binary indicators in the Monitoring Framework. Numbers represent the percentage of elements with each score for coverage. ‘Partially covered’ applies to elements for which the indicator(s) reflect some aspects of the element, but not all. ‘Potentially covered’ applies to elements that could be covered by indicators that are still in development, so there is uncertainty as to whether the final metric(s) produced will adequately cover the element.
